# A high dimensional quantification of mouse defensive behaviours reveals enhanced diversity and stimulus specificity

**DOI:** 10.1101/2020.02.24.961565

**Authors:** Riccardo Storchi, Nina Milosavljevic, Annette E. Allen, Antonio G. Zippo, Aayushi Agnihotri, Timothy F. Cootes, Robert J. Lucas

## Abstract

Instinctive defensive behaviours, consisting of stereotyped sequences of movements and postures, are an essential component of the mouse behavioural repertoire. Since defensive behaviours can be reliably triggered by threatening sensory stimuli, the selection of the most appropriate action depends on the stimulus property. However, since the mouse has a wide repertoire of motor actions, it is not clear which set of movements and postures represent the relevant action. So far this has been empirically identified as a change in locomotion state. However, the extent to which locomotion alone captures the diversity of defensive behaviours and their sensory specificity is unknown.

To tackle this problem we developed a method to obtain a faithful 3D reconstruction of the mouse body that enabled to quantify a wide variety of motor actions. This higher dimensional description revealed that defensive behaviours are more stimulus-specific than indicated by locomotion data. Thus, responses to distinct stimuli that were equivalent in terms of locomotion (e.g. freezing induced by looming and sound) could be discriminated along other dimensions. The enhanced stimulus-specificity was explained by a surprising diversity. A clustering analysis revealed that distinct combinations of movements and postures, giving rise to at least 7 different behaviours, were required to account for stimulus-specificity. Moreover, each stimulus evoked more than one behaviour revealing a robust one-to-many mapping between sensations and behaviours that was not apparent from locomotion data. Our results indicate that diversity and sensory specificity of mouse defensive behaviours unfold in a higher dimensional space spanning multiple motor actions.

## Introduction

Mice are innately able to respond to changes in their sensory landscape by producing sequences of actions aimed at maximizing their welfare and chances for survival. Such spontaneous behaviors as exploration [1, 2], hunting [3, 4], and escape and freeze [5–8], while heterogeneous, share the key property that they can be reproducibly elicited in the lab by controlled sensory stimulation. The ability of sensory stimuli to evoke a reproducible behavioural response in these paradigms makes them an important experimental tool to understand how inputs are encoded and interpreted in the brain, and appropriate actions selected [5, 8–10].

Realizing the full power of this approach, however, relies upon a description of evoked behaviors that is sufficiently complete to encompass the full complexity of the motor responses and to capture the relevant variations across different stimuli or repeated presentations of the same stimulus. Instinctive defensive behaviours, such as escape or freeze have been defined on the basis of a clear phenotype – a sudden change in locomotion state. Thus in the last few years it has been shown that speed, size, luminance and contrast of a looming object have different and predictable effects on locomotion [5–8]. Nevertheless, mice do more than run, and a variety of other body movements as well as changes in body orientation and posture could, at least in principle, contribute to defensive behaviours. In line with this possibility a wider set of defensive behaviours including startle reactions and defensive postures in rearing positions have been qualitatively described in rats [11, 12]. However, until now, a lack of tools to objectively measure types of movement other than locomotion has left that possibility unexplored.

We set out here to ask whether a richer quantification of mouse defensive behaviours was possible and, if so, whether this could provide additional information about the relationship between sensation and actions. To this end we developed a method that enables to obtain a 3D reconstruction of mouse poses. We then used this method to generate a higher dimensional representation of mouse defensive behaviours which enabled to quantify a wide range of body movements and postures.

We found that defensive responses to simple visual and auditory stimuli encompass numerous motor actions and accounting for all those actions provides a richer description of behaviour by increasing the dimensionality of behavioural representation. This increase provides an improved understanding of defensive behaviours in several respects. First, behavioural responses are more specific to distinct stimuli than is apparent simply by measuring locomotion. Second, higher specificity can be explained by the appearance of a richer repertoire of behaviours, with equivalent locomotor responses found to differ in other behavioural dimensions. Third, each class of sensory stimuli can evoke more than one type of behaviour, revealing a robust ‘one-to-many’ map between stimulus and response that is not apparent from locomotion measurements.

## Results

### A method for quantifying multiple motor actions

The first aim of this study was to develop a method that enables to obtain a 3D reconstruction of mouse poses. Five different landmarks on the mouse body (nose tip, left & right ears, neck base and tail base, **Fig. 1A**) were tracked using four cameras mounted at the top of an open field arena that we used throughput the study (**Fig. S1A&B**). The 3D pose of the animal was first reconstructed by triangulation of landmark coordinates across the four camera views (**Fig. 1B, Raw;** see **STAR Methods** section **Reconstruction of 3D poses** and **Fig. S1C-F** for details). This initial reconstruction was then refined by using a method we established for this study (**Fig. 1B, Refined;** see **STAR Methods** section **Reconstruction of 3D poses, Fig. S2** and **Supplementary Movie 1** for details). These pre-processing stages allowed us to describe, on a frame-by-frame basis, the mouse pose ***X*** as

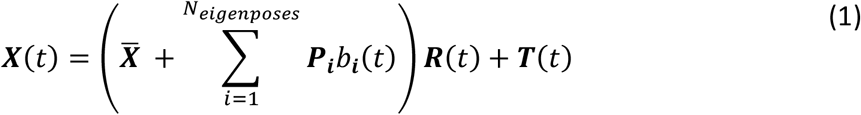

**Figure 1:**
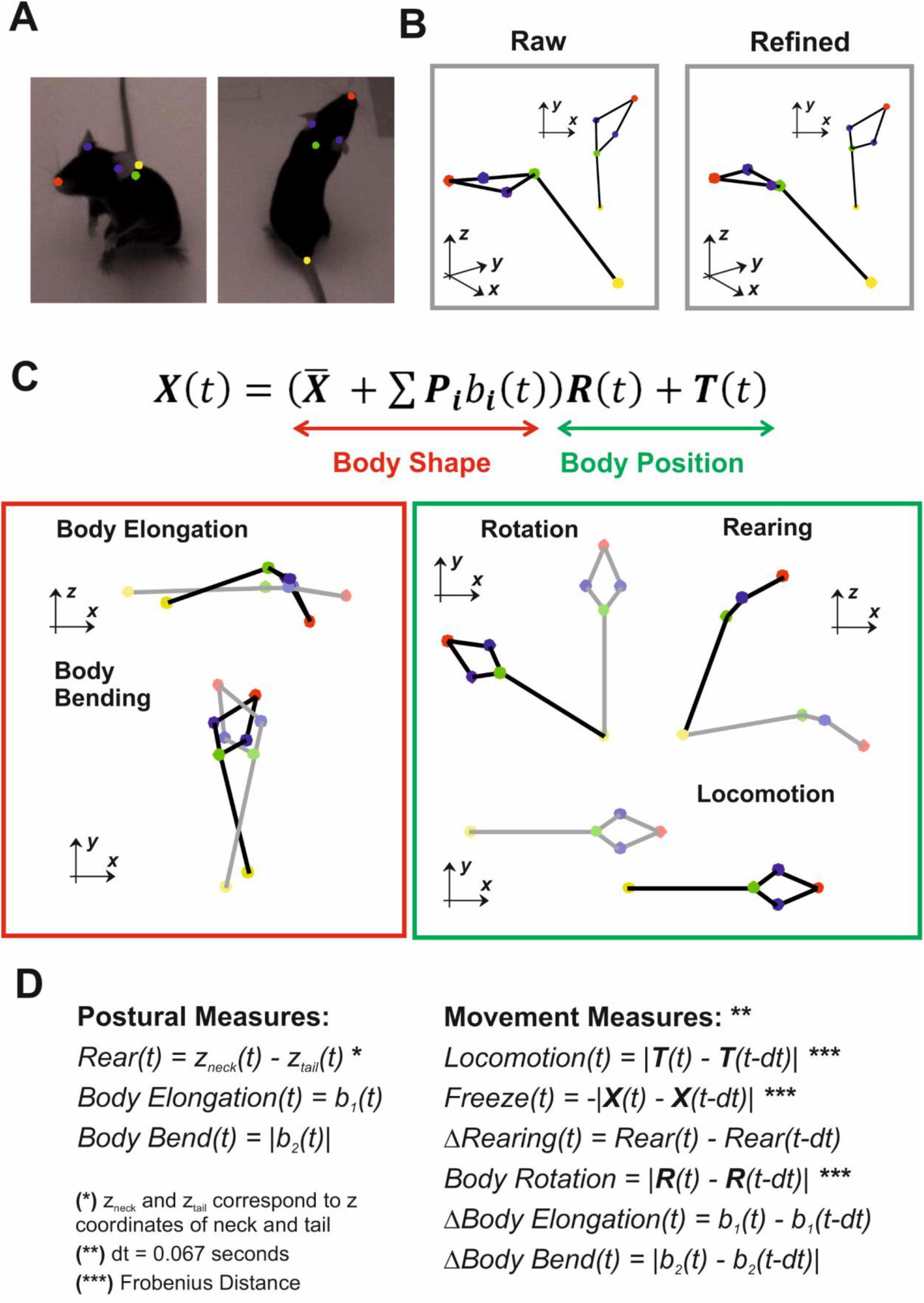
Reconstruction of mouse poses and quantification of postures and movements. **A)** Body landmarks are separately tracked across each camera. **B)** A raw 3D reconstruction is obtained by triangulation of body landmark positions (left panel). The raw reconstruction is corrected by applying our algorithm based on the Statistical Shape Model as described in Methods. The refined 3D reconstruction (right panel) is then used for all the further analyses. **C)** The model expressed by equation 1 allows for quantifying a wide range of postures and movements of which red and green boxes report some examples. The “Body Shape” components enable to measure changes in body shape such as body elongation and body bending. The “Body Position” components enables to quantify translations and rotations in a 3D space. **D)** The full set of behavioural measures, divided into 3 postural measures and 6 movement measures is expressed as function of the terms in equation 1.

Where: *t* represents the time of the current frame; ***X*** the coordinates of the body landmarks; 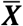 the body coordinates of the mean pose; ***P**_i_* the mouse eigenposes; *b_**i**_* the shape parameters allowing to keep track of the changes in the body shape (**Fig. 1C, Body Shape**); ***R*** and ***T*** the rigid transformations (rotation and translation) encoding the animal’s position in the behavioural arena (**Fig. 1C, Body Position**). Both 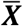 and ***P**_i_* were obtained by training a Statistical Shape Model (SSM, *equation 2* in **STAR Methods** section **Reconstruction of 3D poses**) on a separate dataset of mouse poses. Those poses were first aligned and a principal component analysis was performed to identify the eigenposes ***P**_i_*, i.e. the directions of largest variance with respect to 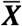. Applying the SSM enabled to correct for outliers in the initial 3D reconstruction and to reduce high dimensional noise while preserving meaningful changes in body shape (see **STAR Methods** section **Validation of the 3D reconstruction** and **Fig. S2** for details). The first two eigenposes captured respectively body elongation and bending (**Fig. 1C, Body Shape**), two important descriptors of the mouse posture that explained respectively 43% and 31% of the variance associated with changes in body shape (see **STAR Methods** section **Interpretation of the eigenposes, Fig. S3** and **Supplementary Movie 2** for details).

Based on this analytical description of the mouse pose we developed two sets of measures to quantify distinct postures and movements. The first set of measures, rearing, body elongation and body bending, allowed us to capture different aspects of the mouse posture (**Fig. 1D, Postural Measures**). The second set, constituted by locomotion, freezing, rigid body rotation and changes in rearing, body elongation and body bending allowed us to capture different types of body movements (**Fig. 1D, Movement Measures**). For all the analyses the measures in **Fig. 1D** were normalized and ranged in the interval [0,1] (see **STAR Methods** section **Normalization of the behavioural measures** for details). These automatic measures were consistent with the human-based identification of walking, body turning, freezing and rearing obtained from manual annotation of the behavioural movies (see **STAR Methods** section **Validation of postural and movement measures** and **Fig. S4** for details).

### Measuring multiple motor actions provides a higher dimensional representation of behaviour

We set out to investigate the extent to which our measures of postures and movements were involved in defensive behaviours. The animals were tested in an open field arena in which no shelter was provided. In order to capture a wide range of behavioural responses we used three different classes of sensory stimuli: two visual, one auditory. Among visual stimuli we selected a bright flash and a looming object. We have previously shown that these two stimuli evoke distinct and opposite behavioural responses, with the former inducing an increase in locomotor activity while the latter abolishes locomotion by inducing freezing behaviour [7]. The auditory stimulus was also previously shown to induce defensive responses such as freeze or startle [6, 13] (see **STAR Methods** sections **Behavioural experiments, Visual and auditory stimuli** and **Experimental set-up** for details on sensory stimuli and experiments).

We separately averaged all trials according to stimulus class and we found that all our measures were involved in defensive behaviours (**Fig. 2A**). To estimate responses divergence (RD) across stimuli we calculated the pairwise Euclidean distance between average responses and we normalized this distance with that obtained by randomizing the association between stimuli and responses (**Fig. 2A, insets;** see **STAR Methods** section **Response Divergence** for details). Across most measures (except rearing for loom and body bend for sound, see **Fig. 2A, insets**) the average response to the flash clearly diverged from those elicited by other stimuli (RD = 6.28±2.40SD, p<0.001 for n = 16 pairwise comparisons, shuffle test). Average responses to looming and sound were all significant but less divergent (**Fig. 2A, insets**; RD = 2.52±1.21SD, p<0.001 for n = 9 pairwise comparisons, shuffle test).

**Figure 2:**
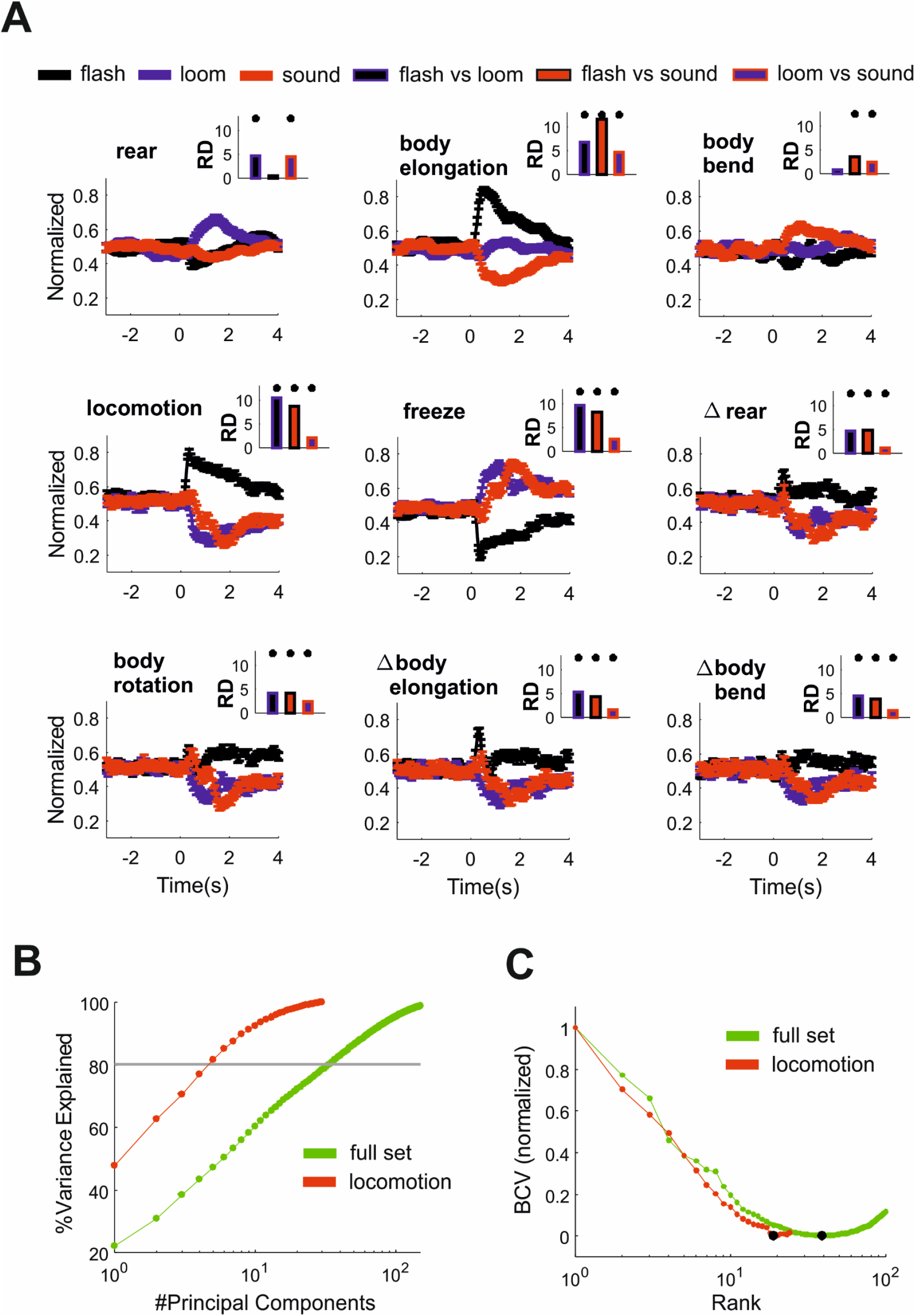
Multiple motor actions are involved in sensory guided behaviours. **A)** Average response to the three classes of sensory stimuli (flash, loom, sound) according to the postural and movement measures defined in **Fig. 1D**. Error bars represent SEM (n=172 for each stimulus class). Response divergence (RD) between pairs of stimuli is reported in insets (*= p<0.001 with shuffle test for RD). **B)** Percentage of variance explained as function of principal components for the full set of motor actions (green) and for locomotion only (red). The grey line indicates 80% explained variance. **C)** The minimum Bi-Cross Validation Error is used to quantify the rank of the full set and of locomotion only (respectively rank = 19 and 39, marked by black dots).

To determine whether the inclusion of all our measures of movements and postures, hereafter the ‘full set’, increased the dimensionality of our behavioural description, we performed a Principal Component Analysis (PCA) on the response matrix. For locomotion, each row of the response matrix represented a trial (n=516 trials) and each trial contained 30 dimensions associated with the 0-2s epoch of the locomotion time series (sample rate = 15 frames/s). For the full set, each trial contained 270 dimensions (30 time points x 9 measures). This analysis revealed that, for the full set, 34 principal components were required to explain >80% variance, while 5 dimensions were sufficient for locomotion alone (**Fig. 2B**). In principle the increase in dimensionality observed in the full set could be trivially explained by a disproportionate increase in measurement noise. To test for this possibility, we estimated the rank of the response matrix by applying the Bi-Cross Validation technique [14] (see **STAR Methods** section **Rank estimation** for details). Consistent with the PCA analysis, we found that the rank of the full set was substantially larger, ~two-fold (**Fig. 2C**), indicating that the full set provided a genuine increase in dimensionality.

### Higher dimensionality reveals increased stimulus specificity in defensive behaviours

We then asked whether this increased dimensionality could capture additional aspects of stimulus-response specificity that could not be observed in locomotion. To account for the fact that evoked responses developed over time we divided the responses into three consecutive epochs of 1s duration according to their latency from the stimulus onset (“early”: 0-1s; “intermediate”: 1-2s; “late”: 2-3s).

We first looked for a specific condition in which the same level of locomotion was expressed in response to two distinct sensory stimuli. A simple illustrative example, where locomotion largely fails to capture stimulus-response specificity, is the case in which both looming and sound induce a common freezing pattern that could be observed in a subset of trials (**Fig. 3A**, top panels; see also **Supplementary Movies 3, 4**). In the intermediate response epoch, when freezing is strongest, locomotion “saturates” towards 0 in responses to both stimuli and thus provides no discrimination (p = 0.48, shuffle test for RD, n = 37 and 31 trials for loom and sound). However, stimulus-specificity is apparent in the animal’s posture as revealed by quantifying body elongation (**Fig. 3A**, bottom panels p = 0.001, shuffle test for RD, n = 37 and 31).

**Figure 3:**
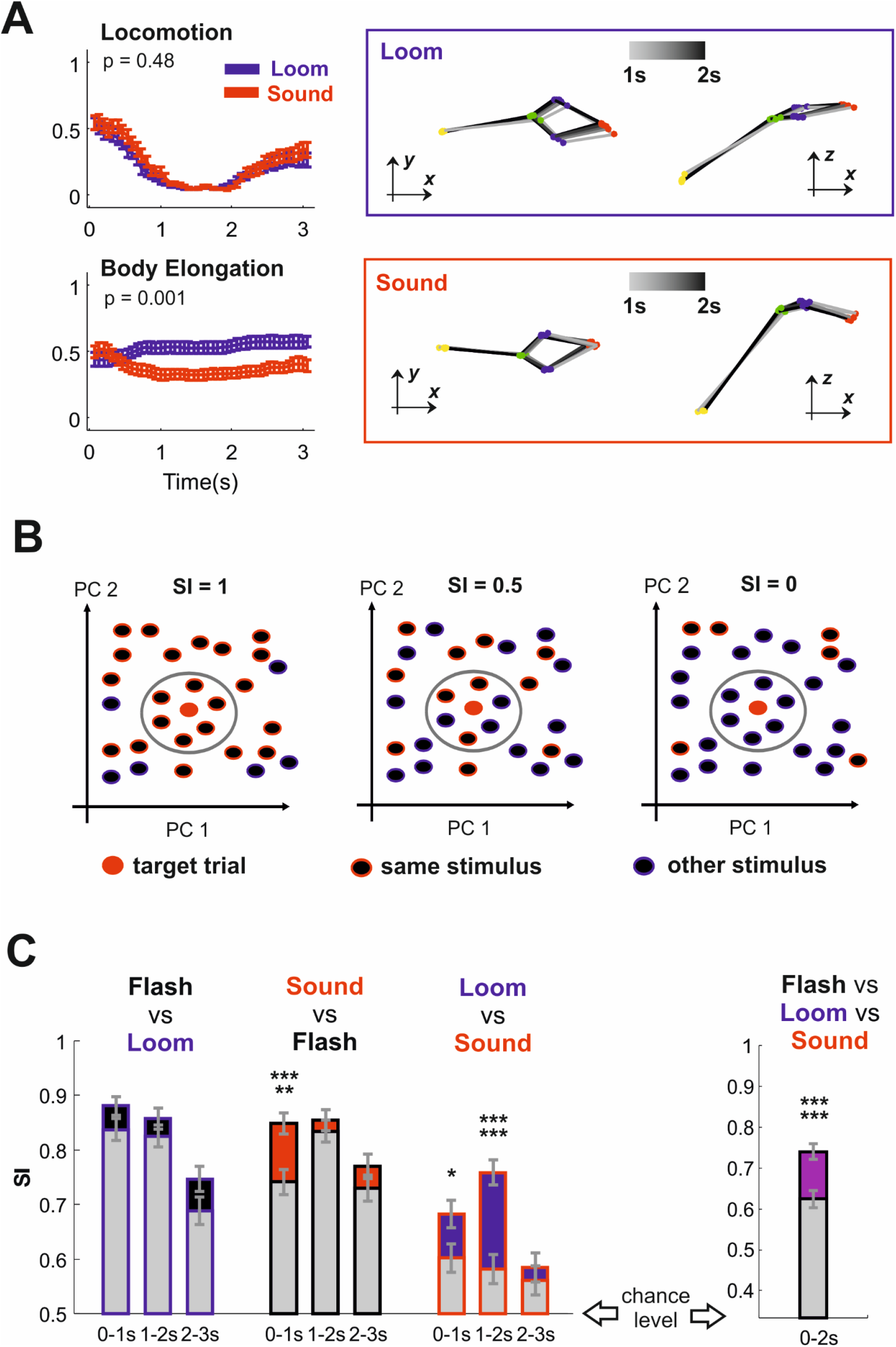
Higher dimensionality reveals increased-stimulus response specificity. **A)** Loom and sound could evoke an indistinguishable pattern of locomotion arrest shown in upper left panel (mean±SEM; data from n = 37 and 31 trials for loom and sound). However the pattern of body elongation was different across loom and sound (bottom left panel). A representative trial for loom (blue box) and for sound (red box) are reported in the right panels. Time progression is captured by the gray-to-black transition of the mouse body (poses sampled every 0.2s between 1 and 2 seconds latency from stimulus onset). Note that different levels of body elongation can be observed from a side view in the z-x planes. **B)** On each trial the specificity index (**SI**) was calculated as the number of neighbour responses to the same stimulus divided by the total number of neighbours. In this toy example, based on two-dimensional responses (PC1 and PC2), we show a target trial for which the number of neighbouring responses for the same stimulus changes across panels to obtain SI values of 1, 0.5 and 0. **C)** Specificity Index for pairs of stimuli (mean±SD, n = 344 trials) measured with locomotion (grey bars) and for the full set (black, red, and blue). **D)** Same as **C** but for all stimuli (mean±SD, n = 516 trials). *p<0.05, ***** p < 0.0005, ****** p < 0.0001.

To systematically compare stimulus-response specificity across all trials (n=172 trials per stimulus) for the full set with the level of specificity revealed by locomotion alone we developed a simple Specificity Index (**SI**). On an individual trial basis, **SI** identified, within a *d*-dimensional space, the *k* most similar behavioural responses across our dataset and quantified the fraction of those responses that were associated with the same stimulus. A toy example in which **SI** is calculated for *k* = 6 in a two dimensional dataset is depicted in **Fig.3B**. Thus, on a given trial, **SI** ranged from 0 to 1, in which 1 signifies all similar behavioural responses being elicited by the same stimulus, 0.5 similar responses being equally expressed for both stimuli, and 0 all similar responses being elicited by another stimulus (**Fig. 3B**). For the real data we used a weighted version of the **SI** index where the contribution of each neighbour response was inversely proportional to its distance from the target response (see **STAR Methods** section **Stimulus-response specificity** for a formal definition of the **SI**). The **SI** was applied to a Principal Component reduction of the response matrix (n = 15 and n = 15 x 9 = 135 time points for locomotion and the full set respectively) and evaluated for pairwise comparisons between the 3 sensory stimuli. Since **SI** was dependent upon *k* and *d* we systematically varied those parameters and we recalculated **SI** for each parameter combination. Almost invariably **SI** was maximized for *k* = 1 both for the full set and for locomotion only (**Fig. S5A**). At least 5 Principal Components were typically required to maximize **SI** and the best value for *d* varied across different comparisons (**Fig. S5B**). Therefore low dimensional responses (e.g. based on the first two components as in **Fig. S5C**) failed to capture the full specificity of behavioural responses. Responses from the same animals were no more similar than those obtained from different animals since, for any given trial in the dataset, the most similar response rarely belonged to the same animal (n = 21, 10 trials out of 516 for full set and locomotion across all stimuli; p = 0.205, 0.957, shuffle test). Moreover the distance between each target trial and its nearest neighbour was on average the same irrespectively of whether they shared the same stimulus or not (**Fig. S5D**).

We then compared **SI** between locomotion and the full set for *k* = 1 and the parameter *d* that returned the highest trial-averaged **SI**. We found no significant differences when comparing flash and loom (**Fig. 3C,** blue bars; p = 0.0674, 0.2416, 0.0701 for 0-1s, 1-2s, 2-3s epochs, sign-test, n = 344 trials). However the full set provided an increase in specificity for early responses when comparing flash and sound (**Fig. 3C,** red bars; p = 0.0002, 0.4570, 0.1980 for 0-1s, 1-2s, 2-3s epochs, sign-test, n = 344 trials) and for the early and intermediate responses when comparing loom and sound (**Fig. 3C,** red bars; p = 0.0254, 0, 0.5935 for 0-1s, 1-2s, 2-3s epochs, sign-test, n = 344 trials). For both the full set and locomotion the highest **SI** values were observed either in the early or intermediate epoch of the response. We then set out to quantify the overall change in specificity. Compared with locomotion, the full set provided an overall ~40% increase in **SI** over chance levels (**Fig. 3D**; p = 0, sign-test, n = 516 trials).

To further test our conclusion that a higher dimensional description of behaviour revealed increased stimulus-response specificity, we asked whether it improved our ability to predict the stimulus class based upon a mouse’s behaviour (i.e. whether higher dimensionality enables more accurate decoding of the stimulus). To this end, we applied a K-Nearest Neighbours (KNN) classifier since this algorithm utilizes that local information provided by the *k* neighbours and therefore represents a natural extension of the specificity analysis (see **STAR Methods** section **Decoding analysis** for details). Like the **SI** index, KNN decoding performances depended on the choice of *k* and *d*. Differently to what we observed for **SI**, where the index was maximized for *k* = 1, the best performances were obtained for larger values of k indicating that multiple neighbours are required to reduce noise (**Fig. S6A**). Similarly to **SI** analyses, high dimensional responses substantially improved accuracy (**Fig. S6B**).

Decoding performances were not significantly different for the full set and for locomotion when comparing flash vs loom (**Fig. 4A**, black dots, p = 0.1130, 0.1384, 0.6013 for 0-1s, 1-2s, 2-3s epochs, binomial test, n = 344 trials). However the full set improved decoding of the early response for flash vs sound (**Fig.4A**, red dots, p = 0, 0.5356, 0.26 for 0-1s, 1-2s, 2-3s epochs, binomial test, n = 344 trials) and across all epochs for loom vs sound (Fig.4A, blue dots, p = 0.0008, 0, 0.0028 for 0-1s, 1-2s, 2-3s epochs, binomial test, n = 344 trials). These results were not specific for the KNN classifier since matching outcomes were obtained by using Random Forest (**Fig. S6C;** flash vs loom: p = 0.4218, 0.2146, 0.3671; flash vs sound: 0, 0.7528, 0.5550; loom vs sound: 0, 0, 0.0057, binomial tests, n = 344 trials). Focussing on the most informative 0-2s epoch enabled to decode flash vs loom and flash vs sound with over 90% accuracy (respectively 93% and 91.73%, **Fig. 4B,** black and red bars) and the full set did not provide significant improvements over locomotion (p = 0.4901, 0.1186, binomial test, n = 516 trials). However, when comparing loom vs sound, locomotion only allowed 66.78% accuracy while the full set provided 77.75%, accuracy, a 65% improvement over chance level (p = 0.00001, binomial test, n = 516 trials). The full set also provided a 20.57% improvement over chance level when decoding was performed across the three stimuli (**Fig. 4C**, purple bar, p = 0.0001, binomial test, n = 516 trials), which corresponded to an additional ~40 correctly decoded trials. Part of the increase in performance was granted by the information provided by changes in body shape (described in **Fig. 1D** as Body Elongation, Body Bend, ΔBody Elongation, ΔBody Bend) since removing those dimensions from the full set significantly degraded decoding performances (**Fig. 4C**, dark purple bar; p = 0.0125, binomial test, n = 516 trials).

**Figure 4:**
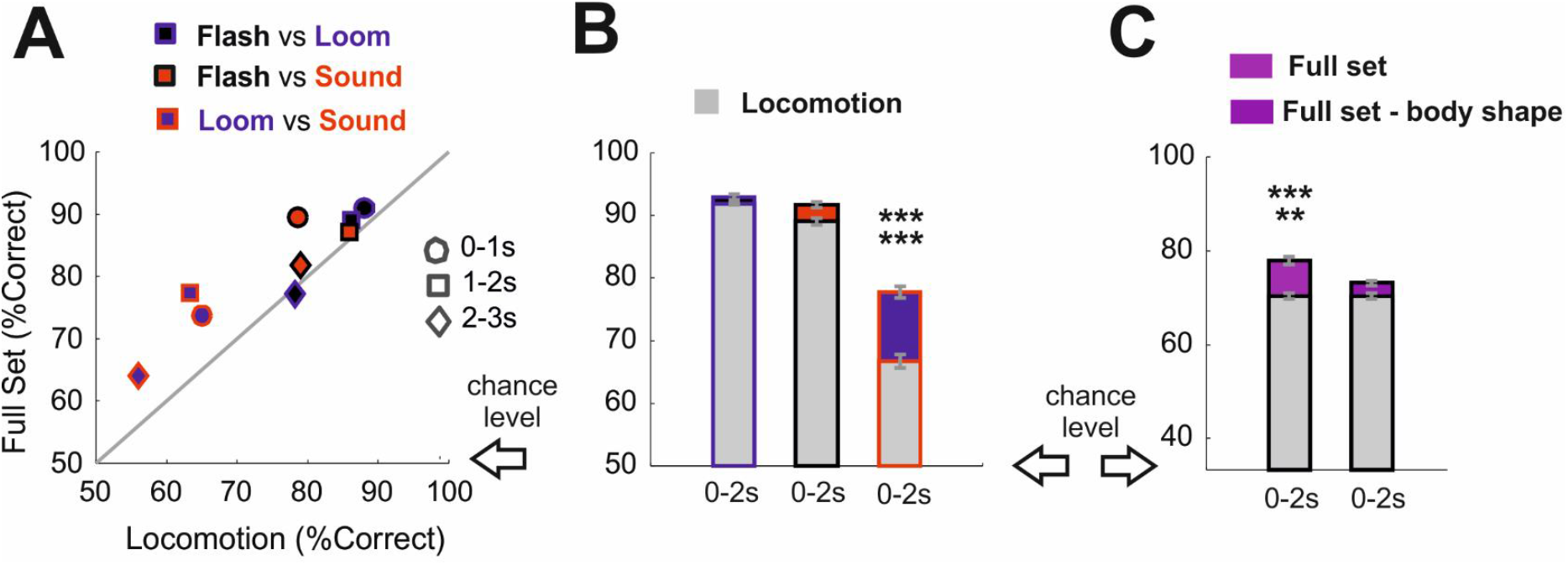
Higher dimensionality improves stimulus decoding. **A)** Comparison between K-Nearest Neighbour (KNN) decoding performances (mean±SD) based on the full set and on locomotion only. Pairwise comparisons are shown for flash vs loom (black-blue), sound vs flash (red-black) and loom vs sound (blue-red) across different response epochs (0-1s, 1-2s, 2-3s). **B)** Decoding performances (mean±SD) of KNN decoding for 0-2s response epochs. **C)** Same as panel **B** but decoding is performed across all stimuli for the full set (bright purple) and for a reduced set in which we removed Body Elongation, Body Bending, ΔBody Elongation and ΔBody Bending (dark purple). Locomotion is always displayed as grey bars. ***** p < 0.0005, ****** p < 0.0001.

### Higher dimensionality reveals a larger set of defensive behaviours

Our results indicate that the mapping between stimulus and behavioural response is more specific in a higher dimensional space. We next sought to describe the structure of this mapping. Specifically, we asked how many distinct behaviours are expressed in response to each stimulus. First, we clustered responses from all trials based upon similarity in motor actions. An important consideration in such a process is how many clusters to allow. We approached that problem by investigating the relationship between the number of clusters and the degree to which each cluster was restricted to a single stimulus (quantified as Mutual Information between stimulus and behavioural response). We focussed on the interval 0-2s since this epoch provided the best decoding results. Then, for each number of clusters, we estimated the Mutual Information (**MI**) between stimulus and behavioural response (see **STAR Methods** section **Clustering and Information Analysis** for details). By observing the increase in **MI** as function of the number of clusters two distinct regions could be clearly delineated (**Fig. 5A,** black error bars). For a small number of clusters, approximately between 2 and 7, we observed a “high gain” region where **MI** increases substantially for each additional cluster. Beyond this domain the “high gain” region was replaced by a “low gain” region where further increments in the number of clusters provided limited increments in **MI**. This analysis suggests 7 clusters as a reasonable trade-off between the need for a generalization of the behavioural responses and the granularity required to capture a large fraction of stimulus specific information.

**Figure 5:**
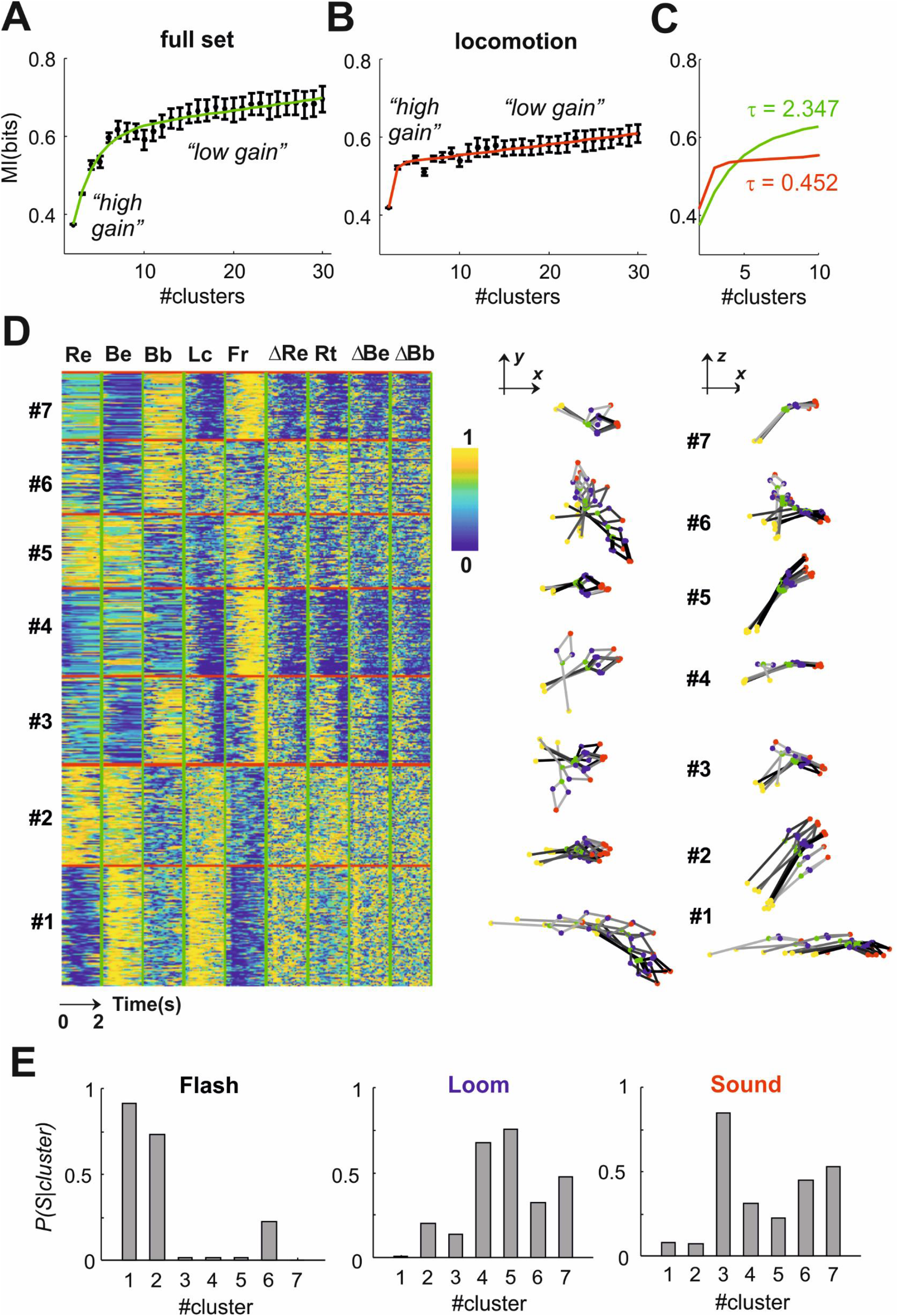
Higher dimensionality reveals a larger set of sensory specific behaviours. **A)** Mutual information is estimated for the full set of motor actions as function of the number of clusters (mean±SD, 50 repeats per cluster; at each repeat the best of 100 runs was selected). Note an initial fast rise in MI (“high gain” region in the plot) followed by a more gradual linear increase (“low gain” region). **B)** Same as panel **A** but for locomotion only. **C)** Comparison between the exponential rise in MI for the full set of motor actions and for locomotion only. The exponential rise in MI, captured by the *τ* values, is slower for the full set indicating that the high gain domain encompasses a larger number of distinct clusters. **D)** Left panel shows the response matrix of the full dataset (n=516 trials) partitioned into 7 clusters. The response matrix is obtained by concatenating all the postures and motor actions (Re = Rear; Be = Body elongation; Bb = Body bend; Lc = Locomotion; Fr = Freeze; ΔRe = ΔRear; Rt = Body rotation; ΔBe = ΔBody elongation; ΔBb = ΔBody bend). Right panels shows one representative trial for each cluster (10 poses sampled at 0.2s intervals between 0 and 2s latency from stimulus onset; time progression is captured by the gray-to-black transition). **E)** Conditional probability of stimulus class, given each of the clusters shown in panel **D**. Flash, Loom and Sound are reported respectively in left, middle and right panel.

Our previous analyses suggested that the range of behaviours is larger when considering the full set vs. locomotion alone (see e.g. **Fig. 3A**). To confirm that this was true, we applied the same clustering method to the locomotion data alone. A similar repartition into high and low gain regions was observed (**Fig. 5B,** black error bars). However, the high gain region domain appeared to be reduced to approximately 2-3 clusters suggesting a reduction in the number of sensory specific behavioural clusters. To more rigorously test whether this was the case we fitted the relation between **MI** and the number of clusters *k* using the function

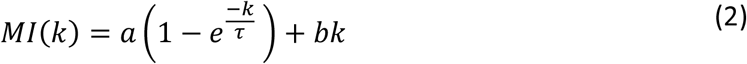

which incorporates a steep exponential component and a more gradual linear component (**Fig. 5A&B**, fitting lines; see **STAR Methods** section **Clustering and Information Analysis** for details). These terms account respectively for the high and the low domain regions. We then used the exponential rise constant ***τ*** as a measure of the size of the high domain region. We found that ***τ*** was indeed smaller for locomotion alone (**Fig. 5C**) indicating that the full set of measures of postures and movements captures a larger number of sensory specific behaviours.

Among the 7 behaviours revealed by our clustering of the full set several motifs occurred (**Fig. 5D**). Fast sustained locomotion (cluster #1) or rearing (cluster #2) both accompanied by body elongation; Body bending followed by delayed freeze (cluster #3); Sustained freeze (cluster #4); Transient freeze in rearing position (cluster #5); Body bending and other rotations of the body axis, including frequent changes in rearing position (cluster #6); Sustained freeze in body bent positions (cluster #7). The Flash stimulus evoked behaviours that were very specific for this stimulus (cluster #1 and #2; **Fig. 5E**, left panel). The Loom and Sound stimuli evoked approximately the same set of behaviours but, between the two stimulus classes, those behaviours were expressed in different proportions (**Fig. 5E**, middle and left panel).

### Distinct behaviours differ both in rate and latency of behavioural primitives

Each of those 7 behaviours was composed of several basic motor actions and postures that we define as primitives. In principle, distinct behaviours could contain diverse sets of primitives and/or the same set of primitives but expressed at different latencies from the stimulus onset. To better understand the composition of each behaviour we increased the temporal resolution of our behavioural analysis by subdividing the 2 seconds window into consecutive sub-second epochs. We then performed a clustering analysis across those sub-second epochs to identify the primitives. In order to select the number of primitives and their duration we used a decoding approach. Thus, for each parameter combination, we fitted three stimulus-specific Variable-order Markov Models (VMMs), one for each stimulus class (see **STAR Methods** section **Analysis of Behavioural Primitives** for details). Decoding performances were then evaluated on hold out data by assigning each trial to the stimulus-specific VMMs associated with the highest likelihood. The VMMs cross-validated performances were optimal for primitive duration between 0.13 and 0.33 seconds (**Fig. S7A**). Within this range the best VMMs contained 6-8 primitives and exhibited maximum Markov order of 0-1 time steps (**Fig. S7B**). We selected VMMs with 8 primitives of 0.13s duration (**Fig. 6A,B**) and we used them to compare, across the 7 behaviours, the rate and the latency of the primitives. For each stimulus the distribution of primitives was significantly different from that observed during the spontaneous behaviour preceding the stimulus (**Fig. S7C;** p = 0, 0, 0, Pearson’s χ^2^ test for flash, loom and sound). For flash the two most frequently occurring primitives defined the responses to cluster #1 and #2 in **Fig. 5D** and represented respectively run and rear actions (**Fig. 6B)**. For loom and sound the most frequent primitives were both expression of freezing but along different postures: with straight elongated body for loom (**Fig.6B, freeze straight**) and with hunched and left or right bent body for sound (**Fig. 6B, freeze bent**). Both the latency and the rate of those primitives changed significantly across the 7 behaviours (**Fig. 6C;** rate: p = 0, 0, 0, 0; latency p = 0, 0, 0.0014, 0; Kruskal-Wallis One-Way ANOVA for run, rear, freeze straight and freeze bent). These results indicate that both the composition and the timing of basic motor actions and postures varies in those behaviours.

**Figure 6:**
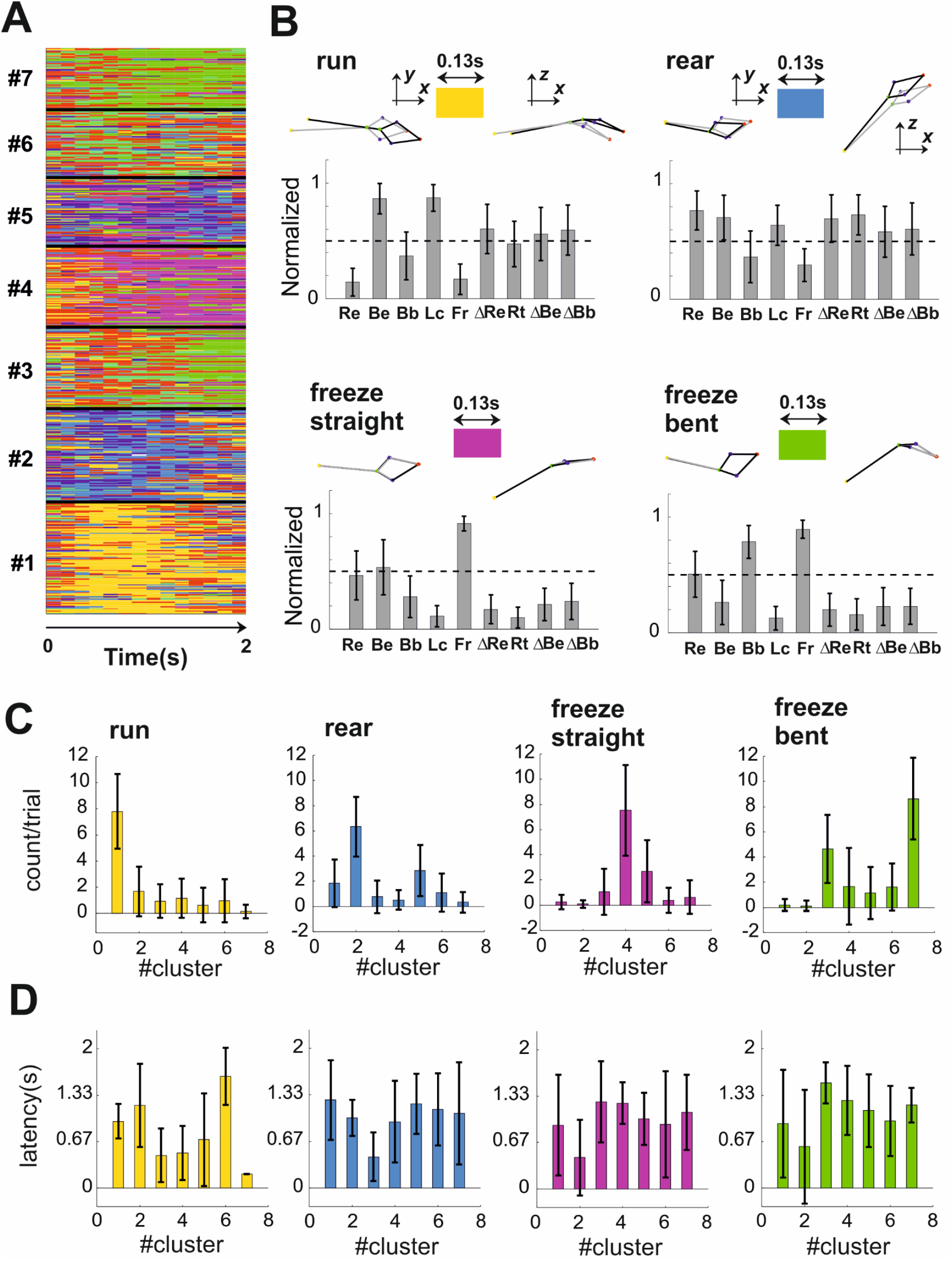
Distinct behaviours differ both in rate and latency of behavioural primitives. **A)** The primitives extracted from the response matrix are displayed for all trials (n = 8 primitives; duration = 0.133s). Trials are partitioned into the 7 clusters as in **Fig.5d**. **B)** The mean±SD of all measures of postures and movements are shown for four primitives (run, rear, freeze straight, freeze bent). Individual representative samples of each primitive are shown as 3D body reconstructions at the top of each bar graph. **C)** Frequency (mean±SD) of each primitive across the 7 behavioural clusters shown in **Fig. 5D**. **D)** Latency (mean±SD) of each primitive across the 7 behavioural clusters shown in **Fig. 5D.**

### The mapping between stimulus and response is not uniquely defined by observable initial conditions

From the results in **Fig. 5** a clear “one-to-many” mapping emerges in which each stimulus can evoke multiple behavioural responses. Such multiplicity could be driven by several factors preceding the time of the stimulus onset and dynamically reconfiguring the mapping between stimulus and response: internal states of the animal that are independent from the stimuli and ongoing observable behaviours; variable postures and motor states that mechanically constrain the range of possible behavioural responses; variable position of eyes and ears within the behavioural arena that modify the way the same stimulus is perceived across trials.

We first set out to explore the effect of ongoing posture and motor state (hereafter for simplicity referred to as ongoing activity). We tested the hypothesis that, given a particular stimulus, the ongoing activity uniquely defined the subsequent behavioural response. To this end, we first performed a clustering analysis on the epochs immediately preceding stimulus onset (duration = 0.5s). Each cluster identified different ongoing activities and the number of clusters was predefined and equal to 7 in order to match the cardinality of the response clusters (**Fig. 7A**). If ongoing activities were to uniquely define the response we would expect a “one-to-one” mapping. We found this not to be the case. Consistently with the “one-to-many” mapping previously described, each ongoing activity cluster led to multiple responses (**Fig. 7B**). To quantify the dependence of response from ongoing activity we used Mutual Information (**MI**). We found that ongoing activity could only account for a small fraction of the **MI** required to optimally predict the responses (14.92% flash, 7.2% loom, 4.77% sound).

**Figure 7:**
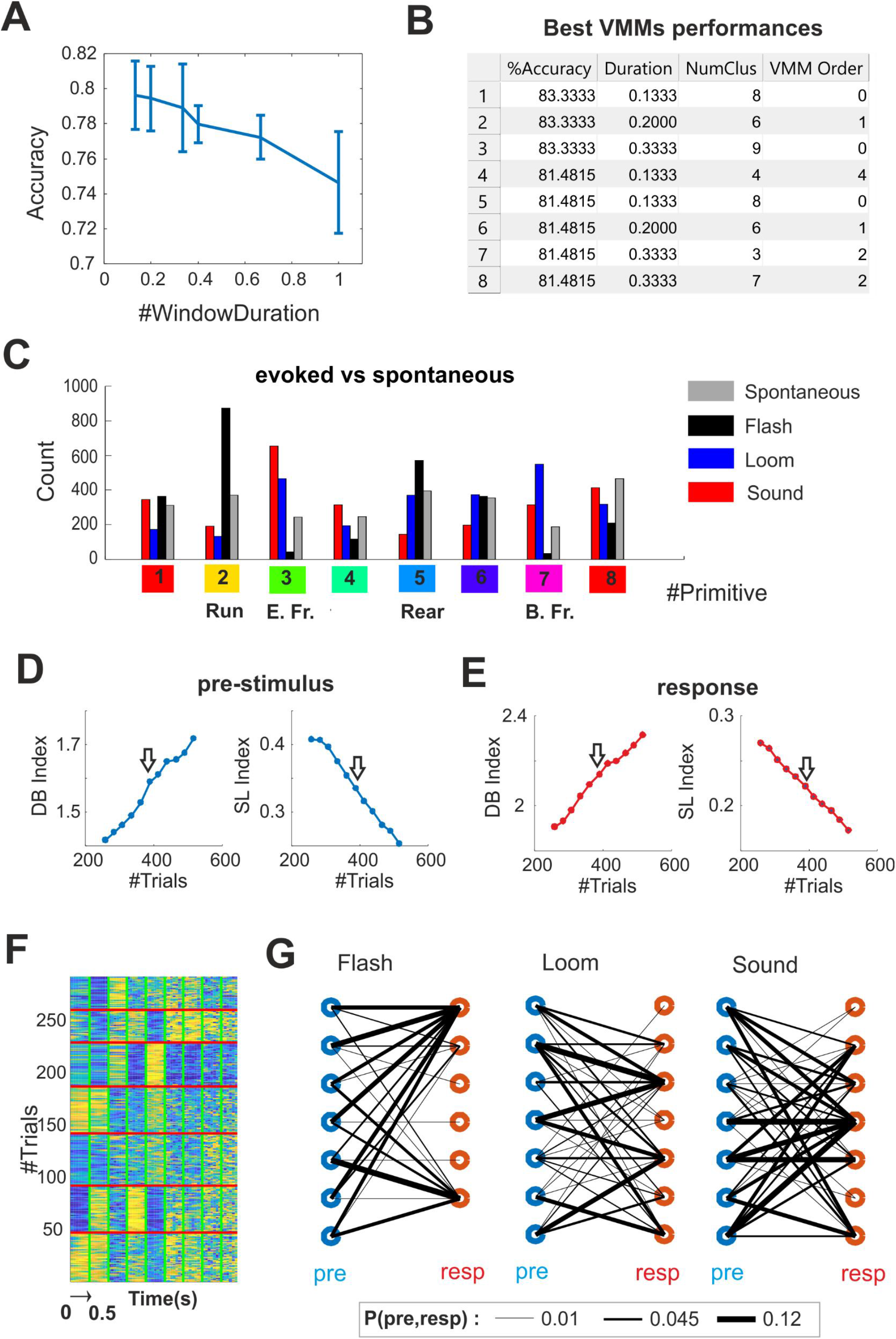
The mapping between stimulus and response is not uniquely defined by initial conditions. **A)** Matrix representing the concatenation of all the measures of posture and movements for the 0.5s preceding the stimulus onset. Trials (n=516) have been partitioned in 7 clusters to match the cardinality of response clustering shown in **Fig. 5D**. **B)** Joint probability of pre-stimulus (blue circles) and response clusters (red circles) for Flash, Loom and Sound stimuli. The probability value is proportional to the width of the lines connecting pre-stimulus and response as shown in legend. **C)** Head elevation is calculated as the vertical angle between nose and neck while head azimuth as the angle of the nose projection on the X-Y plane. **D)** Example of three initial positions. Position 2 is distant from position 1 along the X-Y coordinates but can be exactly superimposed to it by a single rotation along the Z axis. Position 3 is closer to position 1 along the X-Y coordinates but, in order to superimpose these two positions, a translation and two rotations are required. **E)** Example of three partitions of initial positions from the dataset, each pose represents an individual trial. **E)** Mutual Information is estimated as function of the inverse of the overall number of partitions (1/#IP) across 5 dimensions (head elevation and azimuth and head X,Y,Z coordinates). The dotted black lines indicate the entropy of the response clusters.

A caveat of this analysis lies in the fact that the multiplicity of responses might trivially arise from the hard boundaries imposed by the clustering procedure. Thus high dimensional points, representing either ongoing activities or responses, located near the boundaries between two or more clusters would still be assigned to one cluster only. To address the possibility that a “one-to-many” mapping simply arises from trials whose cluster membership is weakly defined we developed a procedure to remove such trials (see **STAR Methods** section **Clustering Refinement**). By removing an increasing number of trials the overall goodness of clustering increased both for ongoing activities and responses (**Fig. S7D,E**). In this reduced dataset (293 trials, out of 516), individual clusters of ongoing activities still led to multiple responses (**Fig. S7F,G**) and only accounted for a small fraction of the **MI** required for correct prediction of the response cluster (11.92% flash, 9.01% loom, 4.85% sound) indicating that the “one-to-many” mapping was robust to clustering errors.

We then set out to investigate the effect of the position of eyes and ears at the time of stimulus onset (hereafter simply referred to as initial position). We quantified initial positions by measuring 5 dimensions: head orientation (elevation and azimuth, **Fig.7C**) and the head X-Y-Z position. All these dimensions were calculated in allocentric coordinates in respect to the centre of the arena (see **STAR Methods** section **Estimating the Effects of Initial Positions**). Since all our measures of movements and postures are instead expressed in egocentric coordinates it is not clear how to connect these two coordinate systems. For example it is possible that initial positions distant from each other in X-Y coordinates but well matched after a rotation around the Z axis would provide more (or less) similar responses than initial positions closer to each other in X-Y coordinates but with poor rotational symmetry (**Fig.7D**). In order to avoid any assumption about the mapping between egocentric responses and allocentric coordinates we developed a systematic method to extrapolate the effect of initial conditions on behavioural responses. Our method relies on the fact that, in the limit of an infinite number of partitions in the space of initial conditions, a “one-to-one” mapping between initial conditions and behavioural responses, if present, will always enable a correct prediction of the response cluster from the initial condition. To test for this possibility we systematically increased the number of partitions (see example partitions in **Fig.7E**) and each time we calculated the **MI** between the initial conditions and the response clusters (see dots black, blue and red dots in **Fig.7F**). We then used linear extrapolation to estimate the **MI** in the limit of an infinite number of partitions clusters (see dots black, blue and red lines in **Fig.7F;** see **STAR Methods** section **Estimating the Effects of Initial Positions**). We found that initial positions only accounted for a minority of the **MI** required for correct prediction of the response cluster (18.76% flash, 10.06% loom, 5.46% sound). Similar results were obtained after removal of 50% of the trials for which the cluster membership for the responses was weakly defined (14.53% flash, 8.46% loom, 8.99% sound). In principle it possible that our linear extrapolation substantially underestimates the information conveyed by initial conditions. However, when the order of the trials for initial conditions and response clusters were separately re-organized to maximize their match, our extrapolation of the **MI** well captured the entropy of the response clusters (92.68%, 95%, 93.03% of entropy for flash, loom and sound; **Fig.7F**, grey dots and lines). This indicates that our extrapolation could capture a “one-to-one” mapping between initial conditions and behavioural responses but such mapping was not present in the data.

## Discussion

A fundamental goal of neuroscience is to link neural circuits to behaviours. Two unescapable tasks are essential prerequisites for approaching this problem: the generation of a detailed anatomical and physiological description of brain circuits – the neural repertoire – and the charting of all the relevant behaviours exhibited by the model organism of choice – the behavioural repertoire. Then, in order to uncover meaningful links, the resolutions of the neural and the behaviour repertoires have to match, since a high resolution on one side can’t compensate for low resolution on the other [15].

In the last decade enormous advances have been made in understanding functional and anatomical connectivity of the CNS [16–18]. Thanks to these techniques a detailed sketching of the neural repertoire underlying sensory guided defensive behaviours in the mouse is in process and substantial advances have been made in the last few years [6, 9, 19–22].

High dimensional reconstruction of rodent behaviour is now starting to catch up (see e.g. [23, 24] for comprehensive reviews). Such reconstructions have been first developed for constrained situations (e.g. treadmill walk) and by applying physical markers to detect body landmarks [25]. More recently, machine learning [26–28] and deep-learning [29–31] have allowed to obviate for the need to use physical markers. Alternative approaches have also been taken by using depth cameras [32] or by combining traditional video with head mounted sensors to measure head movements [33] and even eye movements and pupil constriction [34]. In spite of these advancements, the behavioural repertoire for defensive behaviours has so far only been quantified by measuring changes in locomotion state.

The first aim of this work was to provide a higher resolution map of sensory guided behaviours. To achieve this aim we used four cameras that allowed us to triangulate 2D body landmarks and obtain a 3D reconstruction of the mouse body. The accuracy of such a reconstruction was substantially improved by training 3D Statistical Shape Model that we used to correct the 3D coordinates (**Fig. S2**). Our approach is supervised in that it requires to pre-specify a set of body landmarks (nose, ears, neck base, body centre and tail base; see **Fig. 1A**). Previous approaches to perform a mouse 3D reconstruction, realized by using a depth camera, took instead an unsupervised approach using all body points in the images followed by dimensionality reduction [32, 35]. The main advantage of our supervised approach relies in the fact that the poses are easier to interpret. For example, a mouse looking up can be easily described by a change in nose elevation in respect to the neck base. The main disadvantage is represented by the potential errors in 3D reconstruction arising from incorrect tracking of body landmarks. However, reconstruction errors can be minimized by using multiple camera views and Statistical Shape Models and this approach is easily scalable to any number of views.

Our first main finding was that the level of stimulus-response specificity provided by a high dimensional description of mouse behaviour is higher than the specificity measured with locomotion alone (**Fig.3,4**). This increase in specificity was particularly remarkable when comparing behavioural responses to a loud sound and a visual looming. It has been previously shown that both stimuli induce escape to a shelter or freeze when the shelter is not present [7, 36]. As a result the responses to these stimuli have been considered equivalent and no attempts have been made to differentiate them. Here we show that looming and sound responses can be discriminated with ~78% accuracy (**Fig. 4**). This result can be explained by the fact that a higher dimensional behavioural quantification revealed a larger number of distinct behaviours that are stimulus-specific. Thus for both looming and sound the animals typically froze but they did so according to two different postures: a straight, upward-looking pose for loom (Fig. 3A and cluster #4 in **Fig. 5, 6**) and a hunched pose for sound often preceded by a body spin (Fig. 3A and cluster #3 in **Fig. 5, 6**). Moreover, in several trials a looming stimulus was more likely than sound to elicit rearing or short lasting freeze in rearing position (clusters #2 and #5 in **Fig. 5, 6**).

In locomotion data, where this diversity was lost (**Fig. 5**), specificity for looming and sound was substantially reduced (**Fig. 4**). Linking the neural repertoire to the behavioural repertoire based on locomotion alone would indicate almost perfect convergence – different sensory processes ultimately lead to only one single action. Instead, by increasing the resolution of the behavioural repertoire, we were able to reject the convergence hypothesis showing that behavioural outputs preserve a significant level of stimulus specificity.

For other pairs of stimuli, such as flash vs loom, locomotion alone granted a good level of discrimination (≈90% accuracy, **Fig. 4**). A higher dimensional quantification of postures and movements did not provide substantial advantages in discriminating between such stimuli but enabled to better describe behavioural responses. Therefore, while locomotion data could well differentiate a response to a flash as opposed to a looming stimulus, a higher dimensional quantification could tell us whether the animal was rearing or running (clusters #1 and #2 in **Fig. 5,6**).

Our second main finding was a “one-to-many” mapping between stimulus and response. Thus a high dimensional description revealed at least seven behavioural responses and each stimulus could evoke at least three (**Fig. 5**). The same analysis on locomotion data identified only two behaviours across all stimuli (**Fig. 5B&C**). The reduced, essentially binary, mapping between stimulus and response is consistent with previous results that employed locomotion as unique behavioural descriptor. In absence of shelter a looming stimulation was shown to evoke either immediate freeze or escape followed by freeze [9]. When a shelter was present a dark sweeping object typically evoked a freeze but flight was also observed in a smaller number of trials [5]. Our higher dimensional descriptors provide a substantially enhanced picture of this phenomenon and indicate that the one-to-many mapping between stimulus and response occurs robustly across different sensory stimuli.

The overall figure of seven distinct behaviours represents a conservative estimate and reflects the criterion we used to define the granularity of our behavioural classification. Previous studies, aimed at providing an exhaustive description of spontaneous behaviours, identified of ~60 distinct classes in the mouse [32] and ~100 in fruit-fly [28]. The smaller set of behaviours identified in this study, although more tractable and still sufficient for capturing stimulus-response specificity, likely underestimates the repertoire of mouse defensive actions.

The “one-to-many” mapping we described could not be trivially explained by different initial conditions, i.e. by the variety of postures and motor states or by the position of eyes and ears at the time of stimulus presentation (**Fig. 7**). This is consistent with recent results in drosophila where ongoing behaviour had statistically significant but not deterministic effects on future behaviours [37] and on responses to optogenetic stimulation of descending neurons [38]. Therefore, at least to some extent, the “one-to-many” mapping reflects stimulus-independent variability in the internal state of the animal that generates diversity in the behavioural output. Variability in the internal states could take many forms ranging from noise in the neuronal encoding of the stimuli along the visual and auditory pathways [39] to fluctuating levels of arousal [40, 41] or anxiety [42] and further studies will be required to discriminate among those contributions. The high level of functional degeneracy in neuronal networks (see e.g. [43–45]) provides the suitable substrate for the observed behavioural diversity. The presence of functional degeneracy is consistent with recent studies reporting that the expression of defensive responses can be affected by activation of multiple neuronal pathways [9, 10, 46–50]. However our current understanding of the anatomical and functional substrates of this diversity is still insufficient and limited to the locomotion phenotype. We believe that further investigations of such substrates, matched with a more detailed description of defensive behaviours, represent an important avenue for future studies.

## Acknowledgements

This study was funded by a David Sainsbury Fellowship from National Centre for Replacement, Refinement and Reduction of Animals in Research (NC3Rs) to R.S. (NC/P001505/1) and by an MRC grant to R.J.L. (MR/N012992/1).

## Author Contributions

Conceptualisation, R.S. and R.J.L.; Methodology, R.S., A.G.Z. and T.F.C.; Formal analysis, R.S., A.A. and A.G.Z.; Investigation, R.S. and N.M.; Writing, R.S., N.M., A.E.A. and R.J.L.; Funding Acquisition, R.S. and R.J.L.

## STAR Methods

### RESOURCE AVAIBILITY

#### Lead Contact

Further information and requests for resources, reagents or raw data should be directed to and will be fulfilled by the Lead Contact, Riccardo Storchi

#### Materials Availability

This study did not generate new unique reagents.

#### Data and Code Availability

Data and source codes are available at https://github.com/RStorchi/HighDimDefenseBehaviours

### EXPERIMENTAL MODEL AND SUBJECT DETAILS

#### Animals

In this study we used C57Bl/6 mice (n = 29, all male) obtained from obtained from the Biological Services facility at University of Manchester. All mice were stored in cages of 3 individuals and were provided with food and water ad libitum. Mice were kept on a 12:12 light dark cycle.

#### Ethical Statement

Experiments were conducted in accordance with the Animals, Scientific Procedures Act of 1986 (United Kingdom) and approved by the University of Manchester ethical review committee.

### METHOD DETAILS

#### Behavioural Experiments

The animals were recorded in a square open field arena (dimensions: 30cm x 30 cm; **Fig. S1A** and **S1B**). Experiments were conducted at Zeitgeber time 6 or 18 (respectively n = 14 and 15 animals). During transfer between the cage and the behavioural arena we used the tube handling procedure instead of tail picking, as prescribed in [51], in order to minimise stress and reduce variability across animals. After transferring to the behavioural arena the animals were allowed 10 minutes to habituate to the environment before starting the experiment. Auditory white noise background at 64 dB(C) and background illumination (4.08*10^10^, 1.65*10^13^, 1.94*10^13^ and 2.96*10^13^ photon/cm^2^/s respectively S-cone opsin, Melanopsin, Rhodopsin and M-cone opsin) were delivered throughout habituation and testing. In each experiment we delivered 6 blocks of stimuli where each block was constituted by a flash, a looming and a sound. The order of the stimuli was independently randomised within each block. The inter-stimulus-interval was fixed at 70 seconds.

#### Visual and Auditory Stimuli

The flash stimulus provided diffuse excitation of all photoreceptors (S-cone opsin: 4.43*10^12^ photon/cm^2^/s; Melanopsin: 2.49*10^15^ photon/cm^2^/s; Rhodopsin: 1.98*10^15^ photon/cm^2^/s; M-cone opsin: 7.09*10^14^ photon/cm^2^/s). As looming stimulus we used two variants: a “standard” black looming (87% Michelson Contrast; looming speed = 66deg/s) and a modified looming where the black disc was replaced by a disc with a grating pattern (Spatial Frequency = 0.068 cycles/degree; Michelson Contrast: 35% for white vs grey, 87% for grey vs black, 94% for white vs black; looming speed = 66deg/s). As auditory stimuli we used either a pure tone (C6 at 102 dB(C)) or a white noise (at 89 dB(C)) both presented for 1 second. The selection of looming and sound variants was randomly generated at each trial.

#### Experimental Set-Up

The animals were recorded with 4 programmable cameras (Chamaleon 3 from Point Grey; frame rate = 15Hz). The camera lenses were covered with infrared cut-on filters (Edmund Optics) and fed with constant infrared light. The experiments were controlled by using Psychopy (version 1.82.01) [52]. Frame acquisition was synchronized with the projected images and across cameras by a common electrical trigger delivered by an Arduino Uno board (arduino.cc) controlled by Psychopy through a serial interface (pyserial). Trigger control was enabled on Chamaleon 3 cameras through FlyCapture2 software (from Point Grey). All movies were encoded as M-JPEG from RGB 1280 (W) x 1040 (H) images. For tracking RGB images were converted to grayscale.

In order to deliver the flash stimulation we used two LEDs mounted inside the arena (model LZ4-00B208, LED engin; controlled by T-Cube drivers, Thorlabs). The auditory stimuli were provided by two speakers positioned outside the arena. Background illumination and the looming stimuli were delivered by a projector onto a rear projection screen mounted at the top of the arena. Calculation of retinal irradiance for each photoreceptor was based on Govardovskii templates [53] and lens correction functions [54].

### QUANTIFICATION AND STATISTICAL ANALYSES

#### Reconstruction of 3D poses

Three dimensional reconstruction of the mouse body was based on simultaneously tracked body landmarks from four the cameras (**Fig. S1A&B**). The four camera system was calibrated using the Direct Linear Transform algorithm [55] before data collection by using Lego^®^ objects of known dimensions (**Fig. S1C-F**). The reconstruction error after triangulation was 0.153 ± 0.0884SD cm. For source codes and a detailed description of the calibration process see online material (https://github.com/RStorchi/HighDimDefenseBehaviours/tree/master/3Dcalibration).

After data collection body landmarks were detected independently for each camera by using DeepLabCut software [29]. We used *n* = 5 body landmarks: the nose-tip, the left and right ears, the neck base and the tail base (as shown **Fig. 1A**). When the likelihood of a landmark was higher than 0.5 the landmark was considered valid. Valid landmarks were then used to estimate the 3D coordinates of the body points using least square triangulation. The result of this initial 3D reconstruction was saved as raw reconstruction (**Fig. 1B, Raw**).

The raw reconstruction contained outlier poses caused by incorrect or missing landmark detections (typically occurring when the relevant body parts were occluded). To correct those outliers we developed a method that automatically identifies correctly reconstructed body points and uses the knowledge of the geometrical relations between all points to re-estimate the incorrectly reconstructed (or missing) points. Knowledge of these geometrical relations was provided by a Statistical Shape Model (SSM).

We first estimated a statistical shape model (SSM) of the mouse body based on *n* = 5 body points [56]. This was achieved by using a set of 400 poses, each represented by a *n* × 3 matrix ***X**_train_* whose correct 3D reconstruction was manually assessed. During manual assessment the coordinate of each body landmark across the four cameras was evaluated by a human observer. When all landmark location (n = 20, 5 landmarks for each of the 4 cameras) were approved the associated 3D pose was labelled as correct. Each training pose ***X**_train_* was then aligned to a reference pose using Partial Procrustes Superimposition (PPS) and the mean pose 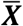 calculated. This algorithm estimates the 3 × 3 rotation matrix ***R*** and the *n* × 3 translation ***T*** matrix that minimize the distance 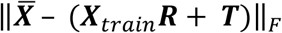 calculated by using the Frobenius norm. A principal component analysis was then performed on the aligned poses to obtain a set of eigenposes **P** and eigenvalues *λ*. The first *p* = 3 eigenposes were sufficient to explain 90.37% of the variance associated with shape changes in our training set (42.68%, 30.85% and 16.84% respectively). Based on those eigenposes the SSM model enabled to express any aligned pose ***X*** as

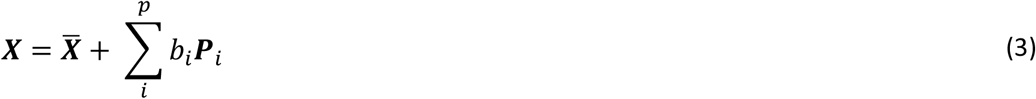

where *b_i_* represent the shape parameters. To identify outlier poses each pose ***X*** was first aligned to the mean pose 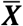 and shape parameters were estimated. A pose was labelled as incorrect when either the Euclidean distance between 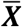 and ***X*** or any of the shape parameters exceeded pre-set thresholds.

Outlier poses could be corrected if only 1-2 body points were incorrectly reconstructed by using the remaining body points and the trained SSM. Correctly reconstructed body points, represented by the (*n* – 2) × 3 matrix ***X**_subset_*, were identified as the subset of points, out of all possible (*n* – 2) subsets, that minimized the distance 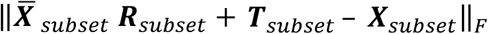. Here the matrices ***R**_subset_* and ***T**_subset_* were obtained by aligning the corresponding body points of the reference pose, 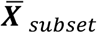, to the selected (*n* – 2) × 3 matrix *X_subset_*. The shape parameters were treated and missing data and re-estimated by applying Piecewise Cubic Hermite Interpolation on the shape parameter time series. The corrected pose ***X*** was then re-estimated as 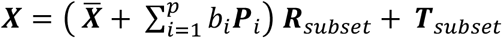.

These preliminary stages enabled to replace gross outliers in the raw 3D reconstruction. We then used all poses ***X*** and associated shape parameters as input for an optimization procedure aimed at obtaining a refined 3D reconstruction by minimizing the following cost function:

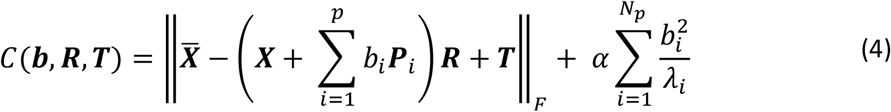

where the right-hand side of the *equation 3* represents a regularization factor to penalize for excessive changes in body shape. The value for the regularization parameter *α*, set at 0.001, was determined by first applying this cost function to a simulated dataset. For all further analyses the time series of each element of ***b, R*** and ***T*** were smoothed using the kernel *w* = [0.2 0.6 0.2]. After smoothing each rotation matrix ***R***(*t*) was renormalized by using Singular Value Decomposition.

Following this reconstruction procedure the mouse pose at any given frame *t* was defined by shape parameters ***b***(*t*) and rigid transformations ***R***(*t*) and ***T***(*t*) as reported in *equation 1*. The final 3D poses were defined as refined reconstruction (**Fig. 1B, Refined**). A dynamic visualization of the refined reconstruction can be found in **Supplementary Movie 1**. All 3D data and source codes for estimating SSM and the refined reconstruction can be found here: https://github.com/RStorchi/HighDimDefenseBehaviours/tree/master/3Dreconstruction

#### Validation of the 3D reconstruction

In order to compare raw and refined poses we first quantified the number of outliers. A pose was defined as outlier when, once aligned with the reference pose, its Euclidean distance from the reference in a 15 dimensional space (5 body points along the X,Y,Z axes) was larger than 5cm. For the raw and refined poses we detected respectively %3.31 (1037/31320) and 1.26% (395/31320) outliers (**Fig. S2A&B**). In the raw 3D reconstruction the outliers were widespread across 178 trials while in the refined 3D reconstruction the outliers were concentrated in 7 trials that were then removed for all the subsequent analyses. Among inlier poses the distance from reference pose was only slightly reduced (**Fig. S2C, inset**). However for the refined inlier poses the distance from the reference pose was fully explained by only 3 components while 9 components were required for the raw inlier poses (**Fig. S2D**). The low dimensional variability associated with the refined inlier poses reflects the constraints imposed by the SSM (via the 3 eigenposes) while the high dimensional variability associated with the raw inlier poses reflects the effect of high dimensional noise. Such low and high dimensional variability can be clearly observed for the whole dataset of inlier poses in **Fig. S2E**.

#### Interpretation of the eigenposes

The SSM enabled to identify a set of eigenposes that captured coordinated changes in the 3D shape of the animal body encompassing all the five body landmarks (see eq.2). To gain more intuitive insights about what type of shape changes were captured by each eigenpose it is useful to visualize those changes. We did so by creating a movie (**Supplementary Movie 2**) where we applied a sinusoidal change to individual shape parameters in *equation 3*. In this way, at any given time t and for the *i^th^* eigenpose, the mouse body could be described as 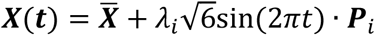. By looking at the movie it is apparent that each eigenpose captures coordinated changes in the distances between body landmarks and angles between head and body. To quantify those changes as function of each eigenpose we selected, based on the movie inspection, a set of four measures: nose-tail distance, neck-tail distance and head-to-body angles on the XY and the YZ planes. We found that the first eigenpose best correlated with nose-tail distance and head-to-body on the YZ plane indicating that this eigenpose captures different levels of body elongation (**Fig. S3A,D**). The second eigenpose best correlated with head-to-body on the XY plane thus capturing left-right bending (**Fig. S3B**). The third eigenpose correlated best with neck-tail distance indicating again a change in body elongation (**Fig. S3C**).

#### Normalization of the behavioural measures

The full set of posture and movement measures was calculated from the refined 3D reconstruction as analytically described in **Fig. 1d**. Each measure was then quantile normalized in the range [0, 1]. First all the values of each measure (n = #time points x #trials = 320 x 516 = 165120) were ranked from low to high. Then, according to its rank, each value was assigned to an interval. Each interval contained the same number of values. The interval containing the lowest values was assigned to 0 and the interval containing the lowest value was assigned to 1. All intermediate intervals were linearly spaced in the range (0,1). Finally the values were converted to their interval number.

#### Validation of the postural and movement measures

In order to validate the measures of postures and movements (**Fig. 1C**) we compared such measures with a manually annotated set. The human observer (AA) watched the behavioural movies and annotated the start and end timing of each action across a subset of data (18 trials from 24 mice, 18 trials/mouse). We focussed on four annotated actions: “Walk”, ‘Turn”, “Freeze” and “Rear”. The action “Turn” included left/right bending of the body as well as full body rotations around its barycentre. The action “Rear” included both climbing up walls and standing on hind legs without touching the walls. All annotated actions lasted on average less than 1 second (“Walk”: 0.71s±0.49s, n = 473; “Turn”: 0.68s±0.42s, n = 214; “Rear”: 0.88s±0.78s, n = 505; mean±SD) except “Freeze” (1.12s±0.70s, n = 371; mean±SD).

Overall the automatic measures of Locomotion, Body Rotation, Freeze and Rearing (**Fig. 1C**) were well matched with manual annotations while also providing additional information about changes in body shape. Thus “Walk” was associated with the largest increase in Locomotion (**Fig. S4A,** right panel) as well as an increase in Body Elongation and decrease in Rearing and Body Bending (**Fig. S4A**, left panel). “Turn” was associated with the largest increase in Body Rotation and Body Bending (**Fig. S4B**). “Freeze” was associated with the largest increase in our measure of Freeze and the largest decrease in Locomotion (**Fig. S4C**, right pane). “Rear” was associated with the largest increase in our measure of Rearing and high sustained Body Elongation (**Fig. S4D**, left pane).

#### Response Divergence

We first calculated the Euclidean distance *D* between average time series obtained from two stimuli. This measure was then normalized by the average distance 〈*D_sh_*〉 obtained by randomly shuffling across trials the association between stimulus and response (n = 1000 shuffles). Finally response divergence was calculated as ((*D* – 〈*D_sh*〉)) / 〈*D_sh*〉. To test for significance we used a shuffle test. We counted the number of times *D* was larger than *D_sh_* and identified response divergence as significant when *D* > *D_sh_* in more than 95% of the shuffle repeats.

#### Rank estimation

For rank estimation we used the Bi-Cross Validation method proposed by (Owen and Perry, 2009). The *m* × *n* response matrix ***X*** is partitioned into four submatrices ***A, B, C, D*** where ***A** ∈ R^r × s^, **B** ∈ R*^*r* × (*n−s*)^, ***C** ∈ R*^(*m−r*) × *s*^, ***A** ∈ R*^(*m−r*) × (*n−s*)^. Then the matrices ***B, C*** and ***D*** could be used to predict ***A***. Specifically if both ***X*** and ***D*** have rank *k* then 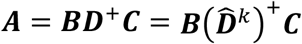 [14], where ***D***^+^ represents the pseudoinverse of ***D*** and 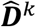 represents the *k*-rank approximation of *D* obtained by Singular Value Decomposition. Using this property we partitioned the rows and columns of ***X*** respectively into *h* and *l* subsets so that each *h × l* subset represented a different hold out matrix ***A***. Finally we estimated the Bi-Cross Validation error as function of the *k*-rank approximation of the ***D*** matrices as:

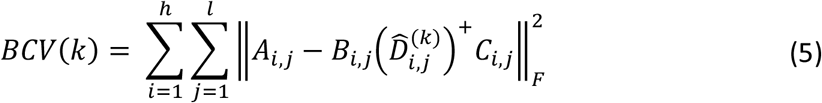

By systematically changing *k* we expect the error would reach its minimum around the true rank of ***X***.

#### Stimulus-response specificity

The Specificity Index (**SI**) for each behavioural response was estimated as the weighted fraction nearest neighbour responses evoked by the same stimulus class. A formal definition of this index is given as follows. Let each *i^th^* behavioural response be quantified by its projection *X_i_* on the *R^d^* space of the first *d* principal components. We define the distance between each pair of responses as *dist_ij_* = ║*X_i_* – *X_j_*║_*L*2_ and its inverse *w_ij_* = 1/*dist_ij_*. The K-neighbourhood of each target response is then defined as the *K* responses associated with the smallest pairwise distances. Let each *i^th^* response be also associated with a variable *Y_i_* = {1, 2} representing the stimulus class. In this way each *i^th^* response is defined by the pair (*X_i_, Y_i_*) ∈ *R^d^* × {1,2}. We can then define *SI_i_*, the Specificity Index for the *i^th^* response as:

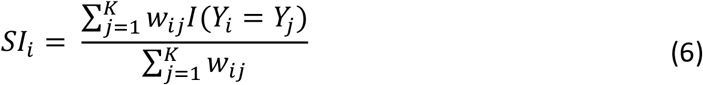

Where the indicator function *I*() is equal to 1 if *Y_i_* = *Y_j_* and 0 otherwise.

#### Decoding Analysis

Decoding performances for K-Nearest Neighbour (KNN) and Random Forest were estimated by using 10-fold cross-validation. Dimensionality reduction based on Principal Component Analysis was performed on the data before training the classifiers. To maximize performances the KNN algorithm was run by systematically varying the parameter K and the number of Principal Components (**Fig. S6A,B**) while the Random Forest algorithm was run by systematically varying the number of Trees (within the set [10, 20, 40, 80, 160, 320]) and the number of Principal Components. Each tree was constrained to express a maximum number of 20 branches. For robustness, the estimates of decoding performances for both KNN and Random Forest were repeated 50 times for each parameter combination. Data and source codes for specificity and decoding analyses can be found here: https://github.com/RStorchi/HighDimDefenseBehaviours/tree/master/Decoding

#### Clustering and Information Analysis

Clustering was performed by using k-means algorithm with k-means++ initialization (Arthur and Vassilvitskii, 2007). The number of clusters k was systematically increased in the range (2-30). For each value of k, clustering was repeated 50 times and for each repeat the best clustering results was selected among 100 independent runs. We then used Shannon’s Mutual Information to estimate the statistical dependence between response clusters and stimuli. A similar approach has been previously applied to neuronal responses (see e.g. [57–59]). In order to estimate Shannon’s Mutual Information the probabilities distributions *p*(*G*) and *p*(*G*|*S*), where 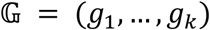 indicates the cluster set and 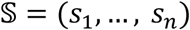 the stimulus set, were estimated directly from the frequency histograms obtained from our dataset. Thus for *p*(*G*) we counted the number of elements in each cluster and we divided by the overall number of elements. We estimated *p*(*G, S*) in the same way and used it to estimate *p*(*G*|*S*) as *p*(*G,S*)/*p*(*S*). From these distributions the response and noise entropies were calculated as

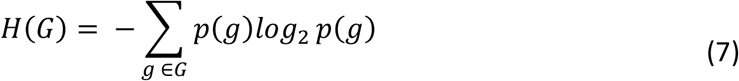

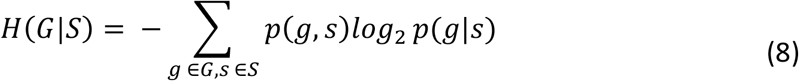

These naïve estimates were then corrected for the sampling bias by using quadratic extrapolation as in [60]. Mutual Information (*MI*) was then calculated from the difference of these corrected estimates. The change in *MI* as function of the number of clusters was fit by using *equation 2* through a mean square error minimization based on the interior point method (Matlab function *fmincon*). For fitting the values of the parameters *a, b* and *τ* were constrained to be positive. Data and source code for clustering analysis can be found here: https://github.com/RStorchi/HighDimDefenseBehaviours/tree/master/Diversity

#### Analysis of Behavioural Primitives

Behavioural primitives were first identified by applying kmeans++ clustering ([61], best of n = 100 replicates for each parameter combination) to the response matrix. For this analysis the response matrix encompassed an epoch starting 0.33s before the stimulus onset and ending 2s after the onset. Since both the number of clusters and the duration of the primitive was unknown we repeated the clustering for a range of [2,10] clusters and for six different durations (0.133s, 0.2s, 0.333s, 0.4s, 0.666s and 1s). In order to model arbitrarily (finite) long temporal relations between subsequent primitives occurring on the same trial we used Variable-order Markov Models (VMMs, [62, 63]). Therefore an additional parameter of this analysis was represented by the maximum Markov order that ranged from 0 (no statistical dependence between two subsequent primitives), to the whole length L of the trial (L = 15, 10, 6, 5, 3 and 2 for primitives of 0.133s, 0.2s, 0.333s, 0.4s, 0.666s and 1s duration). To determine the best VMMs we took a decoding approach. This enabled us to rank the models according to their accuracy in predicting the stimulus on hold-out data. For each combination of cluster cardinality, primitive duration and maximum Markov order we trained three VMMs, one for each stimulus (flash, loom and sound). Thus each of the three VMM (respectively *VMM_fiash_, VMM_loom_, VMM_sound_*) was separately trained by using a lossless compression algorithm based on Prediction by Partial Matching [64] on a subset of trials associated with only one stimulus. On the test set the stimulus 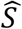 was then decoded by choosing the VMM with highest likelihood 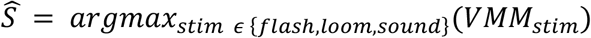. Increasing the temporal resolution of the model by using a larger number of shorter duration primitives increased decoding accuracy (**Fig. S7A**). Parameters for the eight most accurate models are reported in **Fig. S7B.** Data and source code for VMMs analysis can be found here: https://github.com/RStorchi/HighDimDefenseBehaviours/tree/master/VMMs

#### Clustering Refinement

To test the possibility that the “one-to-many” mapping shown in **Fig. 7B** arise from incorrect cluster membership we developed a procedure to improve goodness-of-clustering. The element of each cluster were ranked according to their distance from the centroid. Then for each centroid we removed up to 50% of its elements according to such distance. This resulted in improved clustering metrics as shown in **Fig. S7D&E**.

#### Estimating the Effects of Initial Positions

Initial positions were quantified according to 5 dimensions: head elevation and azimuth, and head X,Y,Z coordinates. In order to partition the space of initial conditions we first generated a set of 5 elements arrays with up to 8 partitions (each partition with the same number of trials) for each dimension (e.g. [1, 3, 4, 1, 1] indicates 3 and 4 partitions respectively along the 2^nd^ and 3^rd^ dimension). For each array in this set the overall number of partitions across the 5 dimensions was the product of the number of partitions in each dimension (e.g. equal to 12 for the previous example). From the initial set we then removed all the items with an overall number of partitions larger than 20. Mutual Information was then estimated for each partition array as described in **Clustering and Information Analysis**. Finally a linear extrapolation was performed to estimate Mutual Information in the limit of an infinite number of partitions.

