## Supplemental Data for "A high dimensional quantification of mouse defensive behaviours reveals enhanced diversity and stimulus specificity"

### Supplemental Information

#### Supplemental Figures

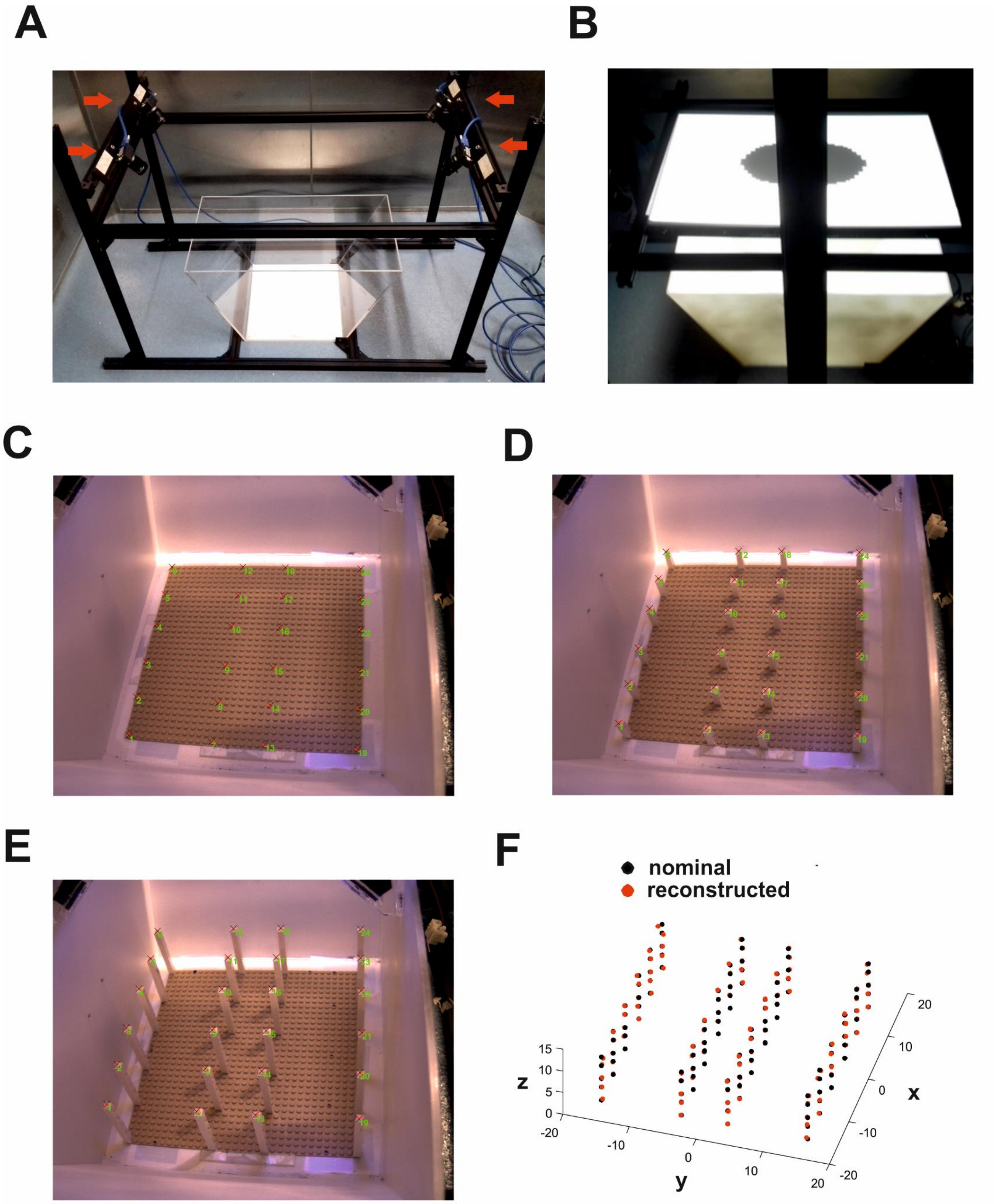

**Figure S1: Calibration of 4 camera system for 3D reconstruction. A)** A picture of our system. The behavioural arena is placed in the centre and cameras are indicated by red arrows. **B)** Visual stimuli, such as the standard black looming disc, are presented through a rear-projection screen. **C)** To calibrate the cameras we used 24 landmarks on a Lego® plate that tiled a large portion of the arena. These landmarks were repeated at 5 different heights by using Lego® bricks as shown in panels **D** and **E. F)** After calibration the 3D reconstructed position of the landmarks (red dots) matched the nominal positions (black dots).

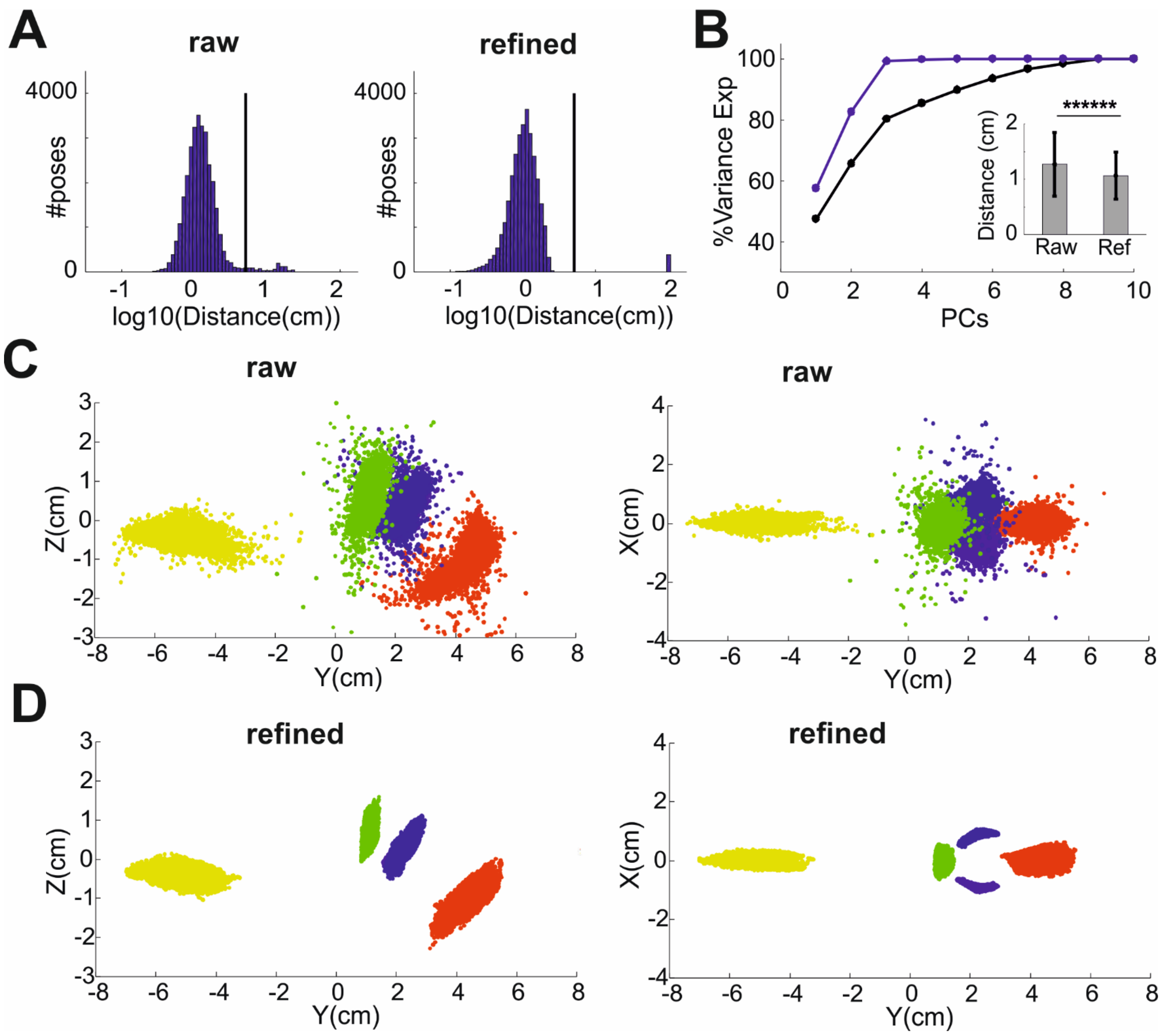


**Figure S2: Validation of the 3D reconstruction. A)** Distribution of Euclidean distances between all poses and the mean pose for raw and refined 3D reconstruction (respectively left and right panel). The threshold that separates outlier and inlier poses is indicated by black vertical lines. B**)** Percentage of variance explained by a Principal Component Analysis of raw and refined 3D poses (respectively blue and black lines). **C)** Full dataset of aligned poses obtained via the raw 3D reconstruction. The 3D poses are shown from a side and top view (respectively left and right panel). **D)** Same as in panel **C** but for the refined 3D reconstruction.

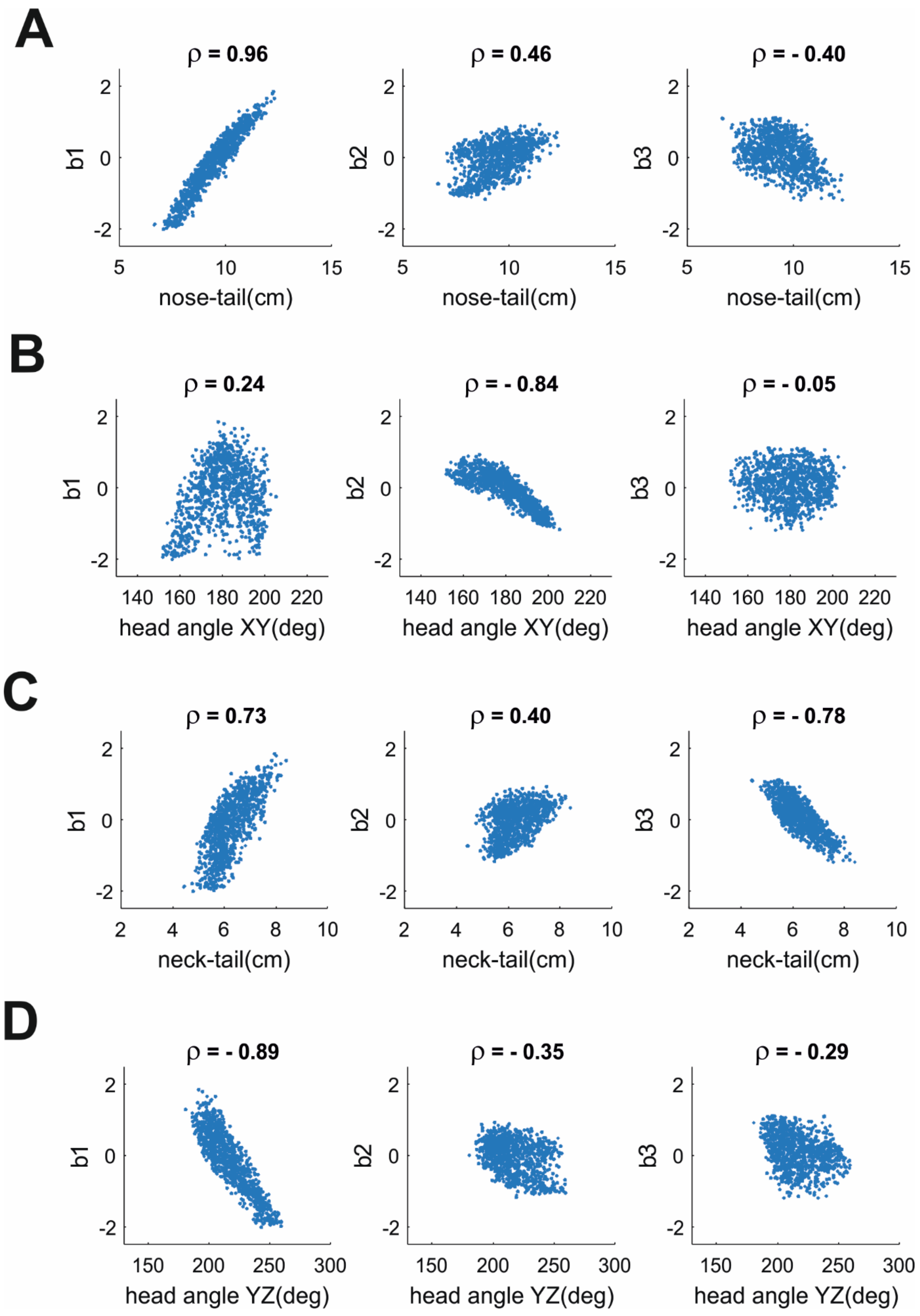


**Figure S3: interpretation of the eigenposes. A)** Shape parameters b1, b2 and b3 are plotted as function of nose-tail distance. **B-D)** Same as panel a but here shape parameters are plotted as function of head-angle on the XY plane (**B**), neck-tail distance (**C**) and head-angle on the YZ plane (**D**). Pearson's correlation values are reported at the top of each panel.

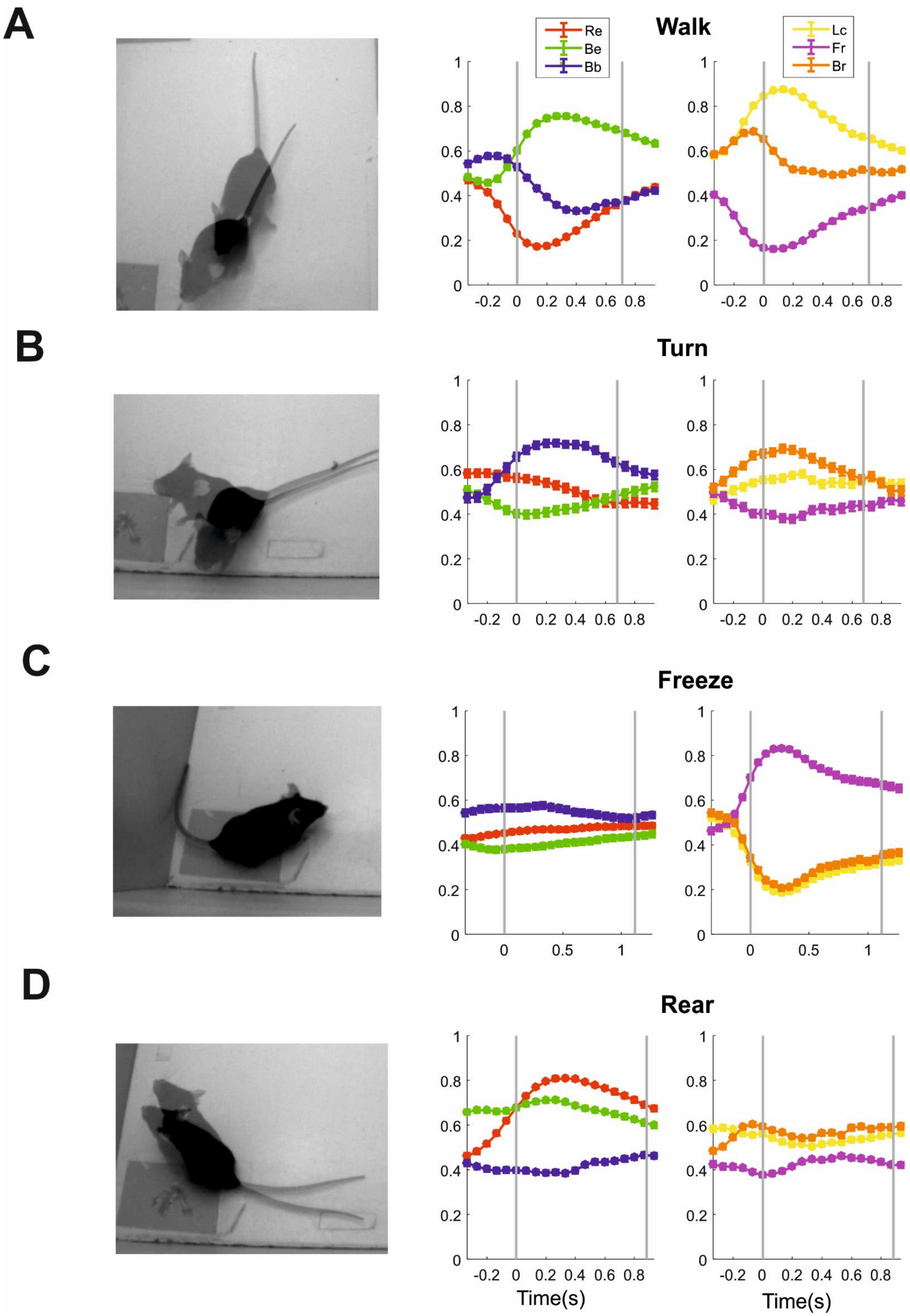

A
Walk
Re
Be
Bb
Lc
Fr
Br
B
Turn
C
Freeze
D
Rear
Time(s)
Time(s)

**Figure S4: Validation of the postural and movement measures.** Manually annotated actions “Walk”, “Turn”, “Freeze” and “Rear” are compared with postural measures of Rearing, Body Elongation and Body Bending (Re, Be and Bb in centre panels) and movement measures of Locomotion, Freezing and Body Rotation (Lc, Fr and Br in left panels). Right panels show representative samples of two superimposed frames 0.267 seconds apart. Centre and left panels show mean±sem of each measure (n = 473, 214, 371 and 505 for Walking, Turning, Freezing and Rearing). In each panel the two vertical grey lines indicate the beginning (at time 0) and average duration of each action.

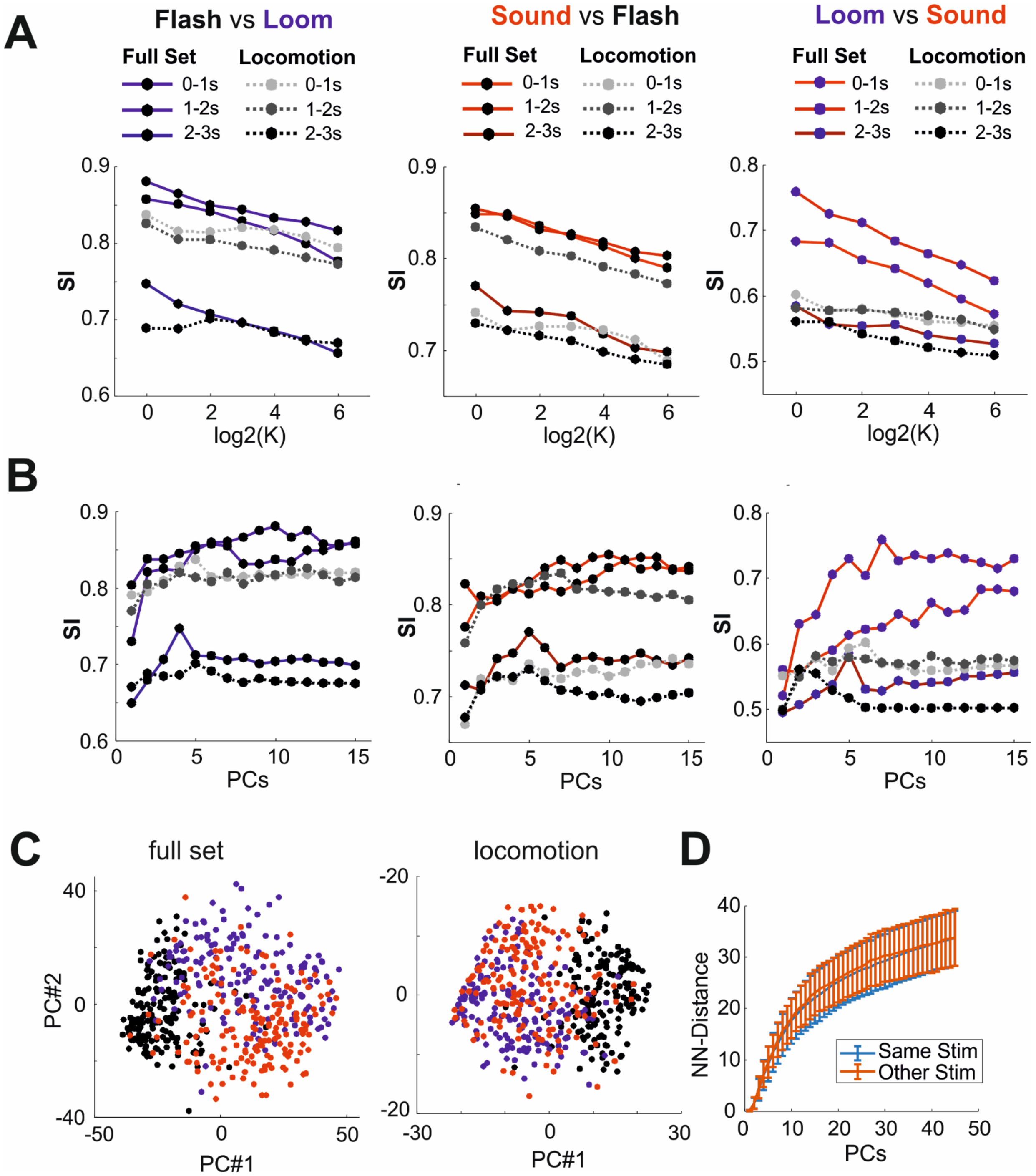


**Figure S5: Higher dimensionality reveals increased specificity in sensory guided behaviours. A)** Specificity Index is shown as function of the number of Principal Components for the full set (continuous lines) and for locomotion only (dashed lines). Different response epochs are represented in different levels of brightness (bright: 0-1s; intermediate: 1-2s; dark: 2-3s). **B)** Same as panel **A** but here SI is shown as function of K. **C)** Scatterplot of the first two Principal Components for full set and locomotion only (respectively left and right panel). Each dot correspond to an individual trial and is coloured according the stimulus (black, blue and red for Flash, Loom and Sound respectively). **D)** Euclidean distance (Mean±SD) of the nearest neighbours to target trials. Blue and Red error bars indicate respectively those trials in which the nearest neighbours was associated with the same (blue) or with a different stimulus (red). None of the 45 pairwise comparisons between the two groups is significant at 0.05 after applying rank-sum tests with Bonferroni correction.

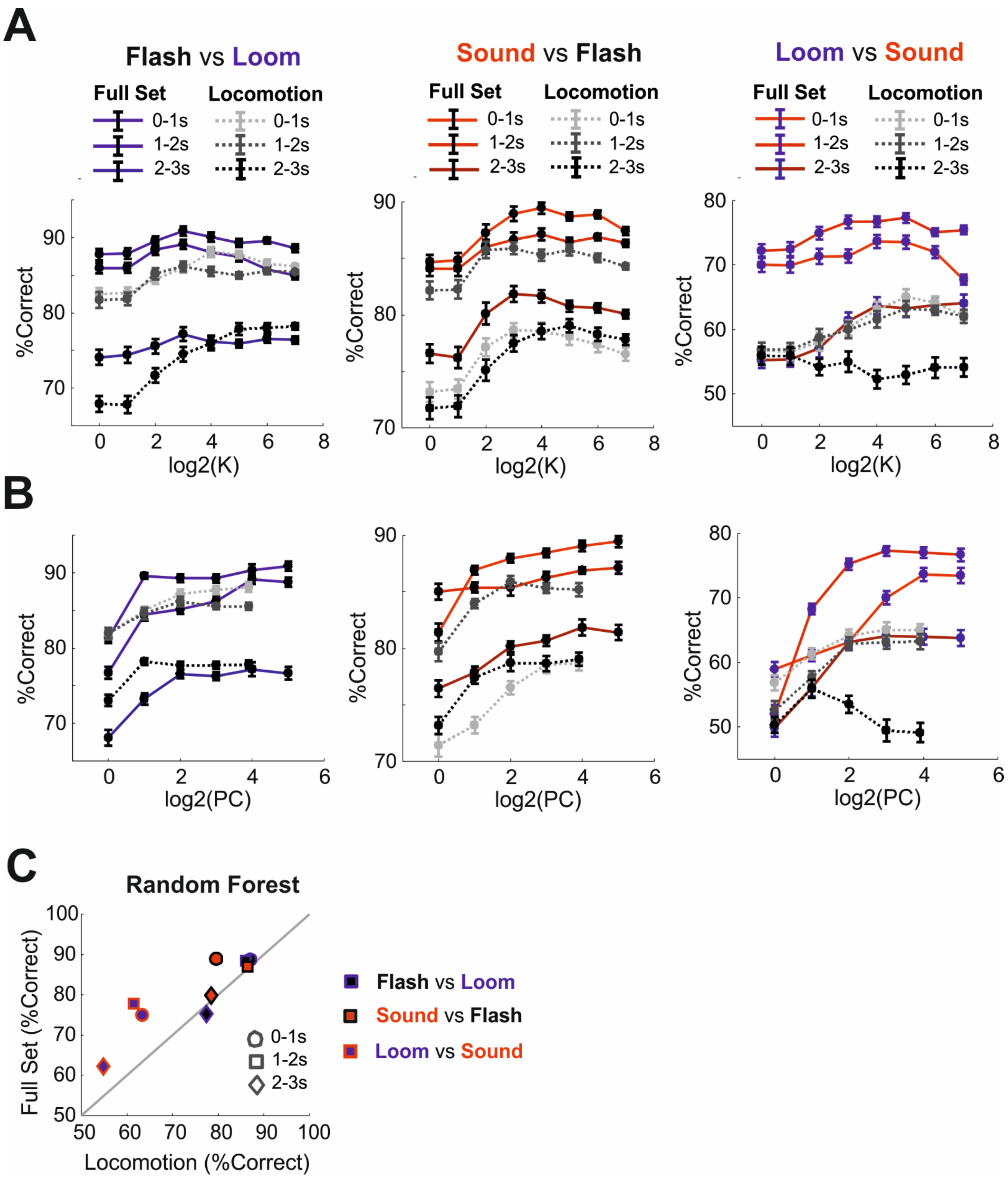


**Figure S6: Higher dimensionality improves stimulus decoding. A)** Stimulus decoding performances across a range of K values for the full set (continuous lines) and for locomotion only (dashed lines). Different response epochs are represented in different levels of brightness (bright: 0-1s; intermediate: 1-2s; dark: 2-3s). **B)** Same as panel a but here the performances are shown as function of the number of Principal Components used for decoding. **C)** Comparison between Random Forest decoding performances based on the full set and on locomotion only. Pairwise comparisons are shown for flash vs loom (black-blue), sound vs flash (red-black) and loom vs sound (blue-red) across different response epochs (0-1s, 1-2s, 2-3s).

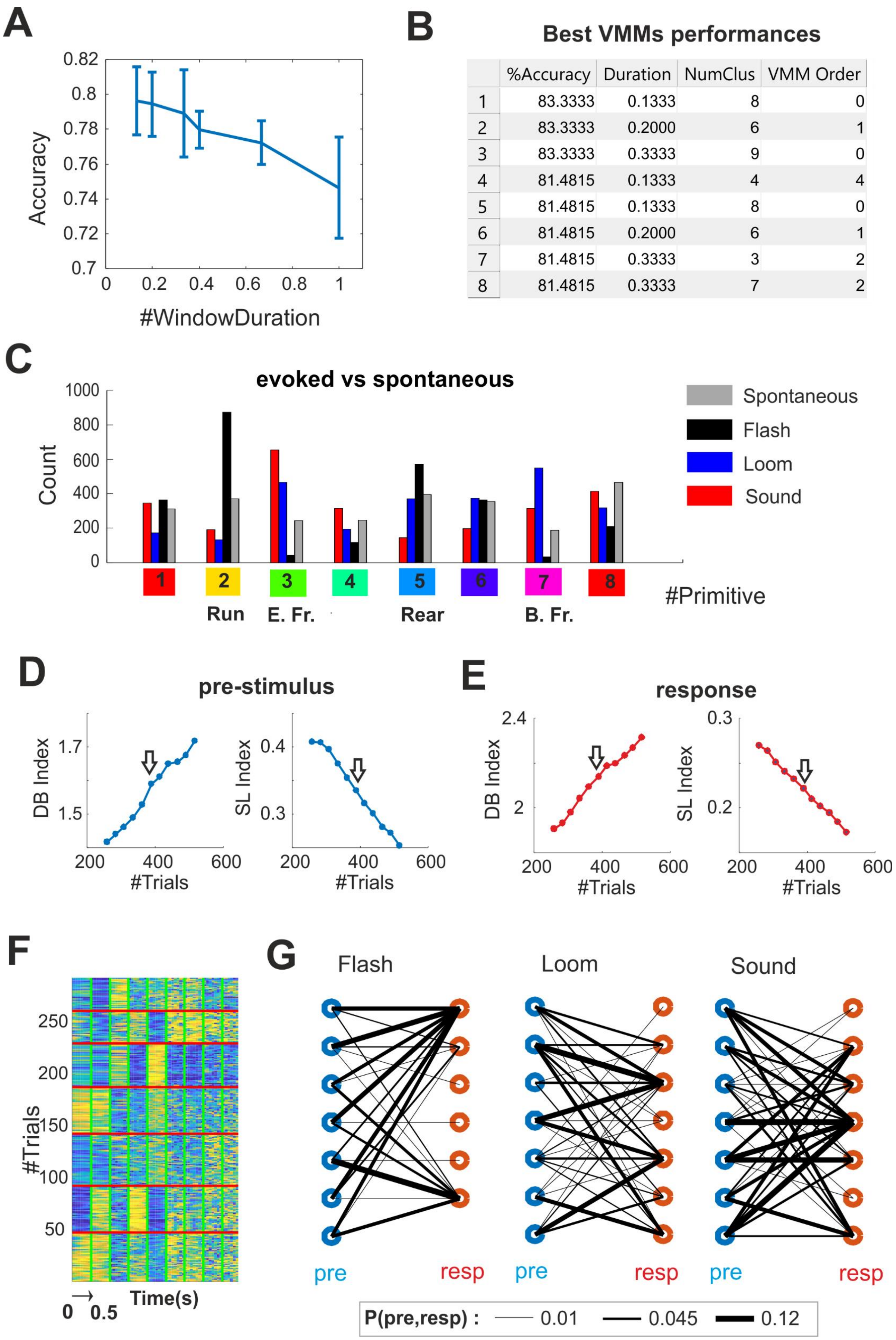


| | %Accuracy | Duration | NumClus | VMM Order |
|---|---|---|---|---|
| 1 | 83.3333 | 0.1333 | 8 | 0 |
| 2 | 83.3333 | 0.2000 | 6 | 1 |
| 3 | 83.3333 | 0.3333 | 9 | 0 |
| 4 | 81.4815 | 0.1333 | 4 | 4 |
| 5 | 81.4815 | 0.1333 | 8 | 0 |
| 6 | 81.4815 | 0.2000 | 6 | 1 |
| 7 | 81.4815 | 0.3333 | 3 | 2 |
| 8 | 81.4815 | 0.3333 | 7 | 2 |

**Figure S7: Distinct behaviours differ both in rate and latency of behavioural primitives. A)** Decoding performances of the VMMs as function of primitive duration (Mean±SD for the 10 best models of each duration). **B)** Defining parameters (Primitive Duration, Number or Clusters, maximum order of VMMs) for the 8 best VOMMs. **C)** Distribution of the 8 primitives for spontaneous activity preceding the stimulus and across responses to flash, loom and sound. **D)** Davies-Bouldin (DB) and Silhouette (SL) indexes as function of the number of pre-stimulus trials. Subset of trials are gradually removed from the full dataset in order to increase goodness-of-clustering as measured by decreasing DB and increasing SL indexes. **E)** Same as panel **D** for response trials. **F,G)** Same as **Fig.7A&B** but here we consider only those trials associated with better DB and SL indexes for pre-stimulus and response (indicated in panels **D** and **E** by arrows).

### Supplemental Movie Legends

**Supplementary Movie 1: Three dimensional reconstruction of mouse poses.** Spontaneous mouse exploration is shown as viewed from two cameras. The refined 3D reconstruction (see **Methods** and right panel of **Figure 1b**) is shown as viewed from the top (top right panel) or from a side (bottom right panel).

**Supplementary Movie 2: Visualization of the eigenposes.** Body shape changes in the directions of the first three eigenposes are animated by sinusoidal modulation of the shape parameters.

**Supplementary Movie 3: Freezing response to a looming stimulus.** A representative trial, displayed as in **Supplementary Movie 1,** shows the behavioural response to a looming stimulus. The time elapsed from the stimulus onset (negative before the onset) is displayed at the top of each panel. The numbers turn red during stimulus presentation.

**Supplementary Movie 4: Freezing response to an auditory stimulus.** Same as in **Supplementary Movie 2** but for an auditory evoked response.
